# ANABAG: Annotated Antibody Antigen dataset with unique features for Antibody Engineering Applications

**DOI:** 10.1101/2025.07.03.663065

**Authors:** Ilyas Grandguillaume, Fernando Luis Barroso da Silva, Catherine Etchebest

**Affiliations:** Université Paris Cité and Université de la Réunion, INSERM, EFS, BIGR U1134, DSIMB Bioinformatics team, F-75015 Paris, France; Université Paris Cité and Université de la Réunion, INSERM, EFS, BIGR U1134, DSIMB Bioinformatics team, F-97715 Saint Denis Messag, France; Programa Interunidades de Pós-Graduação em Bioinformática - USP Av. Prof. Lineu Prestes, 748 Bloco 06 Superior – Sala 667 CEP: 05508-000 – São Paulo-SP – Brasil; University of São Paulo and Université de Paris International Laboratory in Structural Bioinformatics, Faculdade de Ciências Farmacêuticas de Ribeirão Preto, Av. do Café, s/no-Campus da USP, Bloco B, BR-14040-903 Ribeirão Preto,São Paulo, Brazil; Universidade de São Paulo, Departamento de Ciências Biomoleculares, Faculdade de Ciências Farmacêuticas de Ribeirão Preto, Av. Cafe, s/no−Campus da USP, BR, 14040-903 Ribeirao Preto, Sao Paulo, Brazil; Department of Chemical and Biomolecular Engineering, North Carolina State University, Raleigh, NC 27695, United States

## Abstract

The analysis and prediction of antibody–antigen (Ab–Ag) interactions often overlook critical structural features such as glycosylation, physical chemical conditions like pH and salt concentration, as well as the lack of standardized criteria for selecting complexes based on structural properties and sequence identity. Common practices in dataset construction rely on removing redundancy using sequence identity thresholds, which can inadvertently exclude complexes with alternative binding modes that share identical sequences. To enable more precise Ab–Ag modeling and antibody engineering, it is essential to incorporate richer structural and physical information into both physics-based and machine learning models. To address these limitations, we present ANABAG, a new curated dataset of Ab–Ag complexes annotated at the residue level with UniProt sequence information and enriched with a wide range of structural and physicochemical features. The dataset allows flexible filtering of complexes using a variety of descriptors available at both the complex and residue levels. Selected features are ready to use in machine learning workflows, while the structural files are compatible with antibody design and docking pipelines like Rosetta or Haddock. The complete dataset is available on Zenodo, and all accompanying scripts and usage documentation can be accessed via GitHub at https://github.com/DSIMB/anabag-handler.git.

## Introduction

Antibodies are nature’s precision-engineered molecules as part of the immune system response to an exogenic substance, capable of binding to them with specificity [1]. Understanding their interactions is fundamental not only for immunology but also for advancing therapeutic and diagnostic design. When an exogenous substance (i.e., an antigen) enters in contact with a host, a cascade of events is triggered by the innate or adaptive immune system of the host. The differentiated B-cell lymphocytes, called plasmocytes, as part of the adaptive immunity response, produce proteins called antibodies (Ab). These proteins are specialized in the recognition of a specific antigenic-determining region or epitope at the surface of an antigen (Ag) [2]. On the Ab side, the binding region with the antigen is called the paratope. The large number of possible Ags (such as viral proteins, bacterial toxins, allergens, and tumor-associated molecules) leads to an immense diversity in the Ab repertoire [3].

Among the five isotypes of immunoglobulin (Ig) (IgA, IgD, IgE, IgM, and IgG), the blood circulating IgG isotype is the most produced by the organism, studied, and widely applied in medical practices. IgGs are composed of two heavy chains and two light chains, as established by the pioneering works of Porter [4], further advanced by Edelman et al. [5], [6] and completed by Wu, T. T. & Kabat, E. [7]. Since then, Abs are frequently described on the basis of their sequence and structural characteristics. Accordingly, an Ab is formed by the association of the heavy chains composed of three constant (CH) and one variable (VH) domains, and the light chains, composed of one constant (CL) and one variable (VL) domains [2]. From a structural point of view, each of these domains can be classified as an IgG fold [8] and is composed of a β-sandwich of seven or more strands stabilized by disulfide bridges. The loops of an IgG variable domain carry the three Complementary Determining Regions so called CDR 1 to 3, which are mostly responsible for the antigenic recognition [9]. The rest of the Ab is referred to as the Framework Region (noted FR). Each CDR is labelled depending on the chain it belongs to, L1, L2, L3 for the light chain, and H1, H2, H3 for the heavy chain, as shown in Figure 1. The combination of a VH and VL domains of an IgG is called the variable fragment (Fv), and a Fv with the first CH and CL domains is called the antigen-binding fragment (Fab) (see Figure 1). These fragments carry six CDRs and are commonly found in the experimental records e.g. crystallographic structures. The heavy variable domain of camelid heavy chain antibodies (VHH), also called single domain Abs or nanobodies, carries three CDRs and is also commonly found in experimental records (e.g., crystallographic structures). We will refer to Fv, Fab, and VHH as Abs when discussing general properties of antibody-antigen (Ab-Ag) recognition.

**Figure 1:**
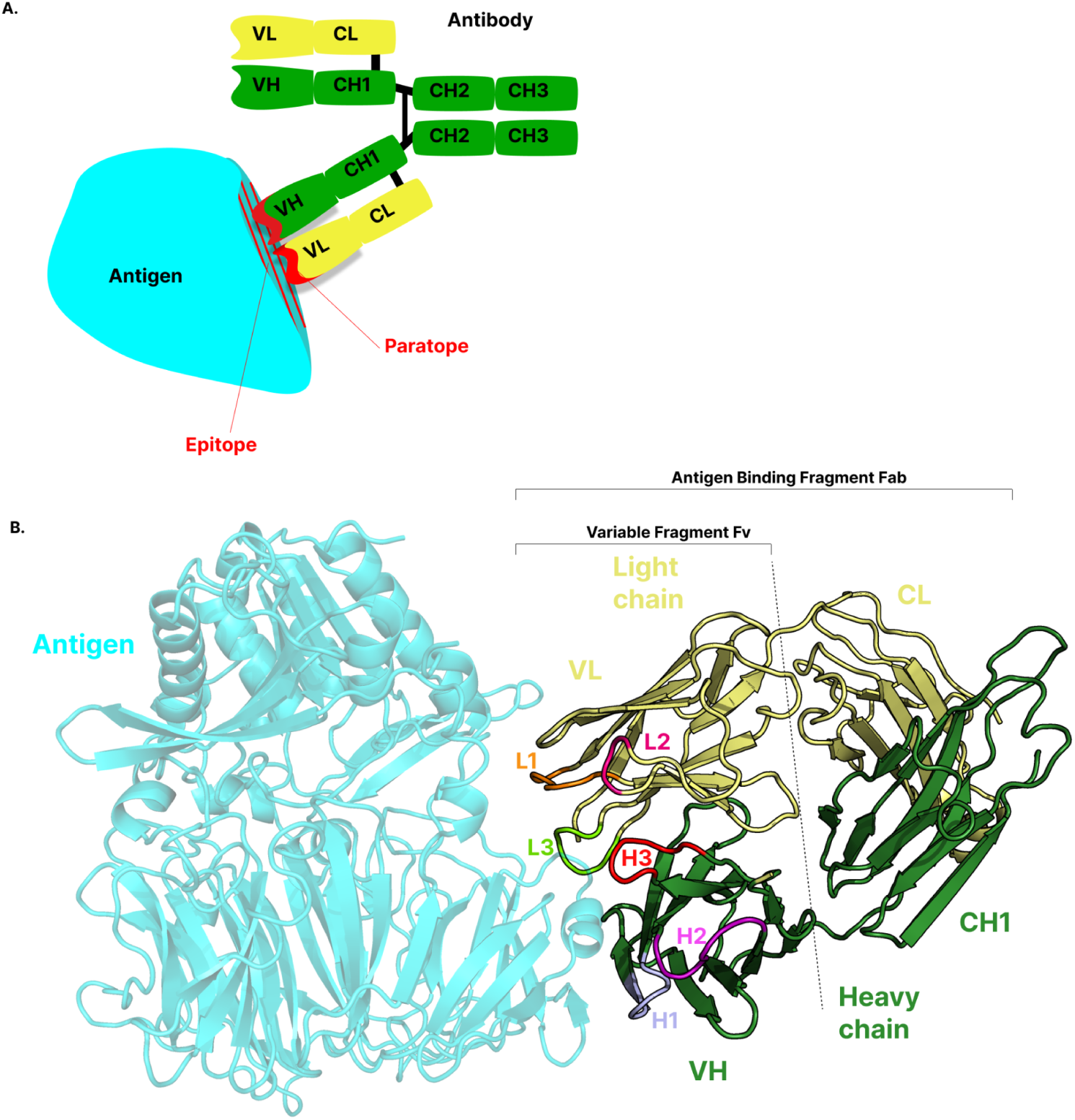
Representations of antibody-antigen (Ag-Ab) complexes and the Ab domains. (A) Top panel: Schematic representation to illustrate the binding interaction between a full-length Ab and its target Ag. The different domains of the heavy (green) and light (yellow) chains are represented as rectangles. The epitope and paratopes are represented in red. (B) Bottom panel: Molecular representation (cartoon) of an Ag-Ab complex (PDB ID: 4FFV). The structure illustrates the molecular interactions between the Ag and Ab, with annotated regions of the Ab, including the Ag-binding (Fab) and the Fv fragments, as well as the complementarity-determining regions (CDRs): L1, L2, L3 (light chain) and H1, H2, H3 (heavy chain).

The expected high specificity of an Ab for a given epitope is attributed to the CDRs. These regions present a large diversity of sequences generated by genetic recombination. This allows a given Ab to recognize specifically an Ag region, while conserving the rest of the sequence and the structure. The amino acids present in each CDR can contribute differently in the Ag-binding process, essentially creating three potential high-affinity regions for different parts of the Ag. Besides the CDRs, the framework region can also play a role in binding, by stabilizing the complex, for instance [10], [11], [12], [13]. It has been shown to be part of the paratope region more often for Ag-VHH complexes [14].

In summary, their natural role in the immune system and their high ability to bind various targets make them attractive candidates for precision-driven design and engineering [15]. They have become indispensable tools in diagnostic but also in therapeutic applications due to their high specificity and decreasing production costs [16]. In this context, accurate prediction of epitope-paratope interactions could significantly improve the efficiency of computational Ab design. Also, it has the potential to reduce part of the experimental efforts to generate high-affinity antibodies during the research and development phase - a process that is often both costly and time-consuming [17].

However, due to their high sequence variability and versatility originating from the combination of three to six CDRs, Abs have been shown to exhibit polyspecificity and play a role in immune and infectious diseases. Polyspecificity refers to the ability of an Ab to bind to another epitope, sometimes in a completely different Ag [18], [19]. This Ab feature leads to difficulties in studying and accurately predicting the epitope-paratope interaction and the potential off-targets of Abs.

Numerous studies have brought insights on the Ab-Ag specific association, and have improved either the paratope, epitope, or interface determination from a sequence or structural point of view [20], [21], [22]. Yet, predicting epitopes from Ag three-dimensional structures remains a difficult challenge, and epitope-intrinsic feature determination remains elusive. This challenge stems from the intrinsic high diversity of Ags and the ability of Abs to bind to a wide variety of surfaces. While numerous studies have successfully identified markers on the paratope - such as an enrichment of tyrosine [1], [23] - the identification of epitope-specific features has proven less straightforward, partly due to the variability in Ag folds and the methods used to analyze them [9], [20], [24].

Although significant progress has been made with deep learning approaches applying Protein Language Models, the success rate is still modest [21], [25], [26]. Two possible explanations might be put forward for this low prediction rate. The first one is the limited availability of data, particularly the valuable structural information from Ab-Ag complexes. A second explanation relies on the choice of the features used for prediction, which is a key aspect, whatever the algorithm used for prediction. Additionally, an important issue that is also frequently underestimated is the impact of the specific physicochemical conditions under which structural information is obtained. This includes the solution pH, the salt concentration, which can further impact the generalizability and accuracy of predictive models.

Concerning the availability of data, among the primary sources of structural information on Ab-Ag are the Protein Data Bank (PDB) [27], the Structural antibody Database (SabDab) [28], and the antibody structure database (AbDb) [28], both of which are built upon the PDB. SabDab, in particular, provides an easy-to-use way of retrieving a great number of Ag-Ab complexes, as well as other useful information about the expression systems, experimental setup, and structural identification of the molecules at play [29].

In the context of studying Ab-Ag interactions, SabDab serves as a classical reference database. It has been widely used to generate datasets for benchmarking predictive methods, such as epitope prediction and docking. Additionally, several datasets have been built upon SabDab, AbDb or the PDB [9], [20], [23], [24], [30], [31], for both benchmark and analysis purposes.

The conclusions drawn from such analysis or the accuracy of the prediction inherently depend on the quantity and quality of available data. Therefore, a thorough examination of the information provided in the PDB is crucial to ensure its reliability, completeness, and applicability. The strategy of creating an Ab-Ag dataset consist in selecting the most representative ensemble of complexes by ***i)*** minimizing the sequence redundancy of Ab sequences by selecting cases that share less than a given identity threshold ***ii)*** minimizing the sequence redundancy of Ag sequences the same way ***iii)*** removing unwanted cases like peptide Ags, and complexes resolved with less than a given resolution (usually 2.5Å).

For instance, Yin & Pierce [32] have built a dataset of approximately 400 complexes to benchmark the efficiency of AlphaFold2.2 in predicting AbAg multimers. The dataset construction was based upon SabDab with stringent criteria on sequence identity between the Ab chains of different complexes, no match between antigen sequence, a threshold for the structural resolution, but also on the structural similarity between components of the complexes. Interestingly, in analyzing the causes of AlphaFold2.2 failures, they highlighted the impact of glycans on predictive accuracy.

Similarly, Xu Z et al. [33] have proposed the pipeline “AbAdapt”, which relies on a dataset of 720 Ab-Ag complexes obtained after applying several filters based on sequence identity, structural similarity, and resolution, as well as the length of the sequence. The dependence between the quantity and quality of data and the method efficiency was clearly stated in their work.

Another benchmark, aiming at evaluating the efficiency of sequence-to-structure prediction methods, was introduced by McCoy et al. [34]. It contains 57 unique Ab–Ag interactions built with a novel method called the “Antibody–Antigen Dataset Maker” (AADaM). This is a small number compared to the larger curated dataset (1,833 complexes) that was proposed by Madsen et al. [24], who applied essentially a resolution threshold on SabDb complexes. A few cases were also removed manually.

Recently, an expanded dataset focusing on Ab structures and docking models has been developed by Almeida et al. [35]. The structures are based on available Abs in the PDB, with a total of 14,184 structures having been identified based on their amino acid sequence. These structures are either free Abs (5,038) or part of a complex (9,146). The crystal packing structures have been kept, and the part of the complexes has been re-docked to constitute an expanded dataset for machine learning.

Among considerations for structure selection, keeping complexes with a large number of missing residues might be problematic and affect the analysis. Yet, discarding the complexes that have a very few number of missing residues might significantly decrease the quantity of data available. Thus, a strategy should be clearly defined to keep a good balance between these two choices. A way consists of building the missing residues, as long as the missing region is relatively short.

Concerning the redundancy, sequence identity thresholds are very useful to remove unwanted redundancy in the dataset. However, since complexes are considered, on which protein(s) the criterion is applied to is a matter of choice. An analysis of the complexes has shown very interesting situations: Ags can be bound by identical or different Abs on distinct epitopes, identical Ags can adopt different conformations, and identical Abs can bind distinct Ags. Importantly, identical complexes can be solved in different physical-chemical experimental conditions.

Consequently, in the present work, we aim at limiting these issues by ***i)*** building a comprehensive dataset gathering information derived from the SabDab dataset on Ag-Ab complexes including VHH, Fab and Fv, ***ii)*** selecting 6,452 biological units (unique set of Ab and Ag chains that form a complex in a crystal structure) over 6,148 initial complexes while assuring maximum diversity by keeping distinct complexes included in the same crystals and removing identical packed complexes, thereby capturing subtle variations that may be critical for understanding binding specificity, ***iii)*** completing most structure gaps in the dataset. Then, once the dataset was built, as a major contribution, we chose to enrich each structure with UniProt annotations but also with a large number of pre-calculated per-residue structural, sequential, and physico-chemical features that we considered to play a key role in the biomolecular interactions. For instance, several unique features incorporated into this dataset are unavailable in other existing datasets: these include the residue charge as a function of pH, the predicted flexibilities of residues, and the detection of glycosylations. This last item further enriches the dataset’s applicability for Ab engineering and the understanding of viral mechanisms.

The dataset presented here, named the ANABAG (ANnotated AntiBody AntiGen) dataset, provides an unprecedented level of functional insight. ANABAG is meant to be a ready-to-use, enriched version of the latest compilation of Ab-Ag structures, with monthly updates. The complexes and features are accessible by downloading them via Zenodo. The full process to access and use the data is described at https://github.com/DSIMB/anabag-handler.git alongside tools to facilitate analysis and structure selection. The huge amount of annotations offers novel features that can be used for prediction purposes or original in-depth analyses of the Ab-Ag complexes. A few examples are also provided in this work.

By addressing challenges related to structural completeness, glycosylation, sequence diversity, and the lack of other critical information, this work provides a framework to enhance computational epitope prediction and, ultimately, antibody design. Additionally, it offers valuable insights for tailoring datasets to optimize predictive models. Furthermore, it enhances the molecular understanding of Ab-Ag interactions by elucidating key physicochemical determinants that govern binding specificity and interaction dynamics.

## Methods

### Dataset Creation

#### Data Extraction

We downloaded the summary file and 3D coordinates of all Ab-Ag complexes from the SabDab database [29] as of 06/01/2025. SabDab is a specialized database for Ab structures based on the PDB. The dataset initially contained 12,175 biological units derived from 6,148 unique PDB entries. In this study, a biological unit refers to a unique set of Ab and Ag chains that form a complex in a crystal structure. For instance, a dimeric Ag interacting with three dimeric Fab fragments in distinct epitope regions would be registered as three distinct biological units. For NMR-derived structures, only the first model was retained. We distinguish the three types of Ab Fragments Fab, Fv, and VHH, based on the length of the Ab sequence. 46 PDB entries that could not be processed because of unidentified residues were discarded, which reduced the number of biological units to 6,102.

##### Filtering of Redundant Biological Units

More than half of the biological units in the dataset were structurally redundant, i.e., the same Ag interacts with the same Ab multiple times, involving the same Ag binding region (often because of the symmetry of oligomeric Ags). This introduces redundancy in the nature of the information that might be extracted. Yet, while many crystal structures contain identical biological units due to Ag symmetry or packing complexes, some exhibit distinct Ag-Ab interactions within the same structure. An example is the Toll-like receptor 3 (PDB id 3ULU), where three Fab fragments bind to different regions of the Ag.

These distinct biological units provide valuable insights into epitope-paratope interactions, which would be lost if only a single biological unit per crystallographic structure is selected, as commonly done in previous studies focused on dataset construction [24]. To preserve all unique Ag-Ab interaction modes, we performed protein sequence alignments using Clustal Omega [36] separately for all Ag and Ab chains given in each crystallographic structure. Sequences that shared 100% identity at non-gap positions were considered redundant. Within each redundant set of sequences, we retained only the corresponding biological unit with the fewest gaps. After filtration, the final dataset contained 6,452 unique biological units from 6,102 distinct PDB structures.

#### Completion

Of the 6,452 biological units in the dataset, 4,237 contained gaps in one or more chains. Since the accuracy of loop modeling decreases with gap length, this information is critical for assessing the reliability of computed structural features. To enhance structural completeness and recover the largest number of functionally relevant segments, we chose to model up to 20 residues to maximize the number of fully completed structures. This resulted in 63% of the incomplete complexes being completed, thus improving the dataset’s applicability for downstream analyses.

The completion was performed using Modeller v10.6 [37], applying its default protocol and parameters for missing regions completion. We did not use AlphaFold [38], since our aim was simply to complete missing regions and not to build the whole structure. Only one model was generated for each biological unit. We did not model gaps located at the N- or C-terminal regions, as these are often unresolved due to experimental factors or added tags.

In total, 241,698 residues among ∼4.69 millions were initially missing across the dataset. We successfully modeled 96,136 of these cases, fully completing 2,926 of the 4,237 complexes that contained missing residues. Note that *all the modelled residues* are clearly stated in the dataset in the *per* residue information, and the total number of modelled residues *per* chain is an available option to select ANABAG complexes.

### Annotation Enrichment

#### UniProt Annotations

To enhance our dataset with important functional information, we mapped UniProt annotations onto the sequences of the 3D structures [39]. This was done by aligning the sequences extracted from the structures with UniProt sequences. Uniprot sequences and annotation were retrieved using the Uniprot browser Retrieve/ID mapping tool [39]. Out of 6,452 biological units, we successfully mapped 5,963 PDB identifiers to their corresponding UniProt entries. We considered only Ags for Uniprot annotations, as Abs were rarely mapped to Uniprot IDs, and when they were, the dissimilarity between the Uniprot Ab sequence and the PDB was important, preventing any meaningful annotation mapping between the two.

For each mapped PDB entry, we retrieved the full UniProt sequence along with the associated annotations and annotation confidence scores. To ensure high alignment quality, we retained only UniProt sequences that shared at least 90% identity with their corresponding PDB-derived sequences. This threshold allowed for minor variations due to differences in protein expression sources while indicating essentially the same protein.

For each pair of aligned sequences, we transferred on the aligned PDB residue, all the UniProt residue annotations, e.g. involved in disulfide bonds, cross-linked residue, belonging to an active or binding site, being glycosylated, lipidated, being a transmembrane residue, involved in a zinc finger or intramembrane region, present in the UniProt sequences that fall within the boundaries of the PDB sequences. For all annotated residues, we also reported their corresponding annotation score.

#### Glycosylation Identification

To identify glycosylation sites in the selected biological units, we used Privateer v. MK V [40], a tool for carbohydrate detection and verification. Through this approach, we detected 15,486 glycosylation sites across all ANABAG structures.

#### Residue Surface accessible Area Calculation

We used Freesasa v. 2.1.2 [40] with the default parameters, based on the Shrake-Rupley algorithm with a probe radius of 1.4 Å, to compute the solvent-accessible surface area (SASA in Å²) of each residue in both the complexed and uncomplexed states. This calculation was used to identify residues that contribute to the interface.

#### Definition of the Interface

Different criteria can be used to define an interface [41]. Interatomic distances are commonly used [20], which means considering the pairs of residues that fall below a chosen threshold to be part of the interface. The other measure that is frequently utilized is the change in the SASA of residues between the bound and unbound forms [42]. The last criterion was chosen here to define the Ab-Ag interface. We calculated the SASA of all residues in the complex and in the isolated molecule. Any residue that experiences a change in SASA between the unbound and bound states is considered to be part of the interface.

### Evaluating Dataset Diversity at the Sequence and Structure Levels

In order to facilitate the use of our dataset for either learning or benchmarking purposes, we carefully analysed the sequence diversity of our dataset obtained using MMseqs2 [42]. Thus, we used MMseqs2 to cluster Ag and Ab sequences at different identity thresholds. The parameters used can be found in the supplementary information (SM, Section Material & Method: How to reproduce the work). For Abs, we applied a 60%, 80%, 95%, and 100% sequence identity threshold. The groups of sequences are respectively referred to as G60AB, G80AB, G95AB, and G100AB (GxAB groupings). Using a 95% identity threshold is a common strategy to cluster antibodies in the field [22], [24], and as they are very similar proteins, there is no need to cluster them with a minimum sequence identity threshold lower than 60% (See results table 3).

For Ags, we used 20%, 40%, 60%, 80%, 95%, and 100% identity thresholds to assess sequence diversity at various evolutionary distances, respectively referred to as G20AG, G40AG, G60AG, G80AG, G95AG, and G100AG (GxAG groupings). This choice, previously used in studies for epitope prediction [21], provides information about conserved and variable residues across Ag sequences. Each GxAG or GxAB clustering is a distinct definition of Ag or Ab sequence redundancy, where sequences in the same cluster or in different clusters are considered, respectively, similar or distinct, all depending on the percent identity used for the clustering.

We performed clustering on all Ag and Ab sequences from the 6,452 biological units using each of the aforementioned identity thresholds. Each sequence was assigned a cluster label at every threshold, creating six distinct groupings for Ags that ranged from diverse (G20AG) to identical (G100AG). For Abs, four groupings were considered. For each cluster, the global multiple sequence alignment (MSA) was obtained using Clustal Omega version 1.2.4 [36], applying default parameters. The alignment indices of each residue were recorded. The alignment indices range from 1 to the length of the MSA and are shared by each residue at a given position.

This allowed us to assess epitope conservation across similar Ags and compare the interfaces of similar Ags and Abs. We evaluated the Ag-Ab pairing redundancy for each cluster of Ag sequence across all GxAG groupings. This means that for a given GxAG, we analysed each cluster obtained with two or more sequences and then evaluated the Ab-Ag interface redundancy of the corresponding complexes. We found that for a given Ag cluster composed of different Ab-Ag complexes, there is often a diversity in interfaces (See the results section in Table 3).

To quantify this redundancy with robust measures, we computed the epitope aligned-index overlap and the amino acid interface similarity between all pairs of complexes within a cluster using the coefficient of interface overlap (CIO) and the Cosine similarity. The process of evaluating the interface redundancy for two complexes having their Ag sequence in the same GxAG cluster is illustrated in Figure 2.

**Figure 2:**
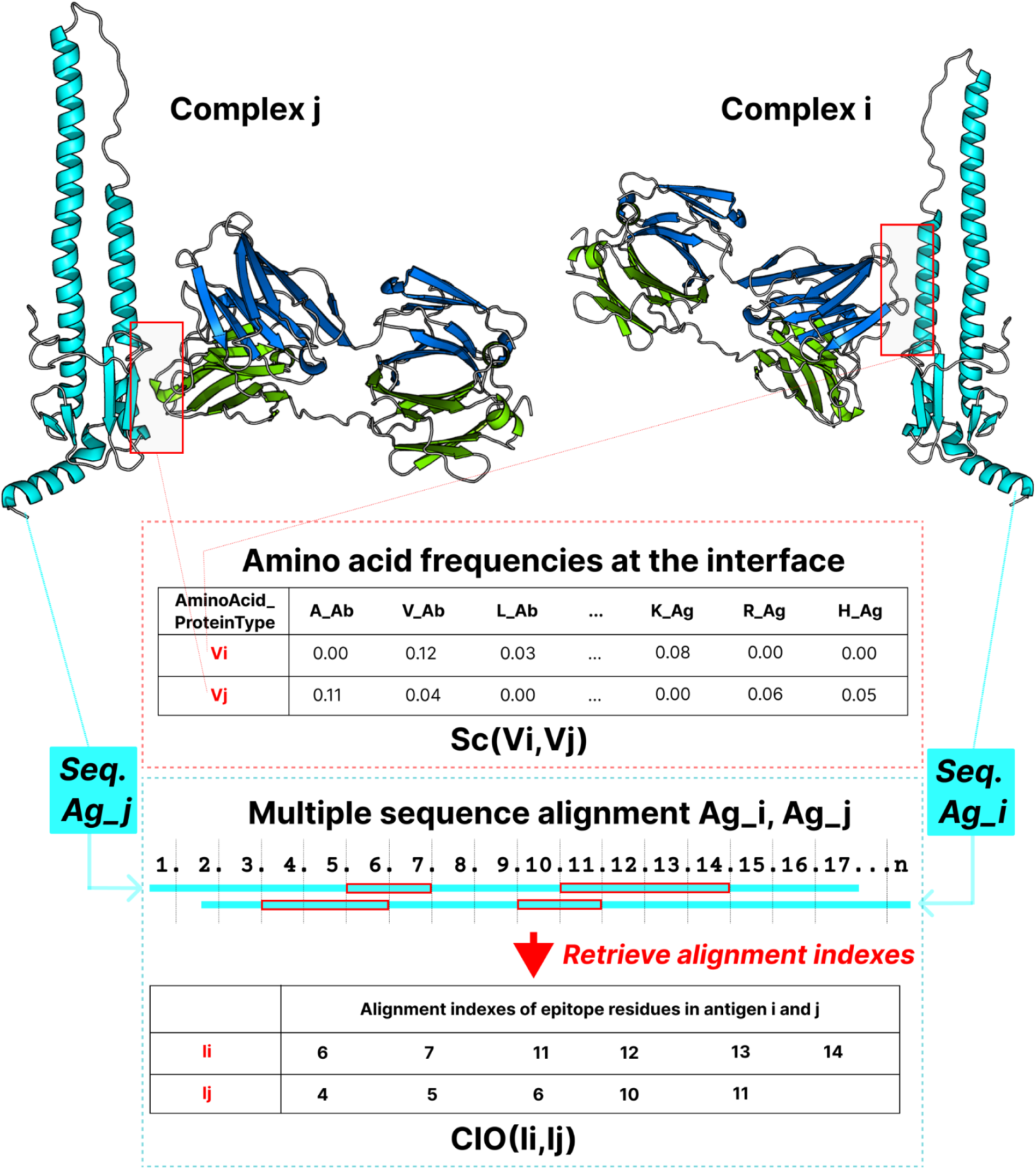
Schematic representation of the procedure used to evaluate interface redundancy between two antigen-antibody complexes (i and j) sharing a common antigen (same GxAG cluster). The amino acid frequencies for each type of protein, e.g., Ab or Ag at their respective interfaces are encoded as frequency vectors (Vi and Vj), and their similarity is assessed via cosine similarity (Sc). A multiple sequence alignment (MSA) of the antigen sequences is also constructed, from which the aligned epitope positions are extracted to compute the coefficient of interface overlap.

The CIO is an adaptation of the Jaccard [43] similarity used to compare the similarity and overlap between two sets of residue indexes as:

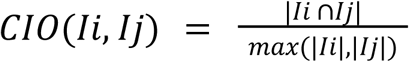

where *I_i_* and *I_j_* are the sets of indexes corresponding to the epitope residues for Ag *i* and *j,* respectively. The indices are derived from the MSA, where the sequences of Ag *i* and *j* are aligned. CIO measures the alignment overlap between two interfaces.

A CIO of 1 signifies that the epitope residues of Ag *i* and *j* correspond to the same positions in the MSA. However, the nature of the amino acid is not fully considered in this measure. To go further, a measure of the similarity of amino acid composition between two compared interfaces was considered, i.e., the cosine similarity [44] between two amino acid frequency vectors. We measure the amino acid similarity between interfaces *i* and *j* with:

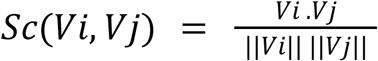

where *V_i_*and *V_j_* are the concatenated amino acid frequency vectors of length 20 for the epitope and paratope for interface *i* and *j*, respectively.

Two interfaces *i* and *j* are considered similar based on both the CIO and Sc using the following condition:

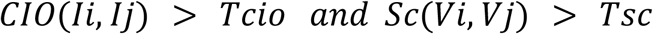

where Tcio and Tsc are the respective thresholds for overlap and amino acid similarity. Tcio and Tcos thresholds have been selected by calculating the Calinski Harabasz index (CH) [45], which is a commonly used measure to select a threshold for unsupervised clustering that is based on intra and inter cluster variance:

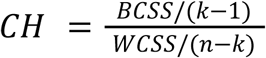

where *k* is the number of clusters and *n* is the number of interfaces. The Between-Cluster Sum of Squares (BCSS) is the weighted sum of squared Euclidean distances between each cluster average and the overall data average [45]. It can be interpreted as the inter-cluster variance. The greater the BCSS, the more distinctive the clusters are.

The Within-Cluster Sum of Squares (WCSS) is the weighted sum of squared Euclidean distances between each data point and its respective cluster average [45]. It represents the intra-cluster variance. The lower WCSS, the more similar the data points within a cluster are. The larger the CH index, the more distinct the clusters and the more compact the data points within a cluster. We selected Tcio = 0.9 and Tcos = 0.9 as the combination of these two thresholds provided clusters of interfaces that maximised the CH index.

## Results

### Dataset characteristics

Starting from Sabdab, a total of 12,175 biological units derived from 6,148 unique PDB entries were curated by applying the filtering criteria detailed in the Methods section. This curation has led to 6,452 unique biological units from 6,102 unique PDB entries composing the ANABAG dataset. The difference between the number of PDB entries and the final number of complexes is due to the existence of distinct Ab-Ag pairings within the same PDB entry.

The average reported resolution for complexes in the dataset is 2.93Å a maximum of 30 Å for electron microscopy, a minimum of 0.96Å, and a standard deviation of 1.64Å (see Figure S.2). The average pH used in experiments to solve the complexes of this dataset is 7.14, with a standard deviation equal to 1.06 (see Figure S3). Importantly, we have noted that some structures have been solved with rather extreme values of pH, i.e., a solution pH of 3.3 for the most acidic case and 11 for the most basic one. How these values could impact the Ab-Ag binding is a key question that we considered by calculating the net charges of the titratable residues as a function of pH using the FPTS method (see below and the SM for a detailed description of the method and parameters used).

#### Dataset annotation and calculated features

As a result of our annotation pipeline, we successfully mapped 64,642 residue-level annotations and 140,544 region-level annotations derived from UniProt (see Table S1). In addition to these mappings, we systematically computed a broad set of structural, sequential, and physicochemical features *for each residue* in every complex. An overview of all included features is presented in Table 1.

**Table 1:**
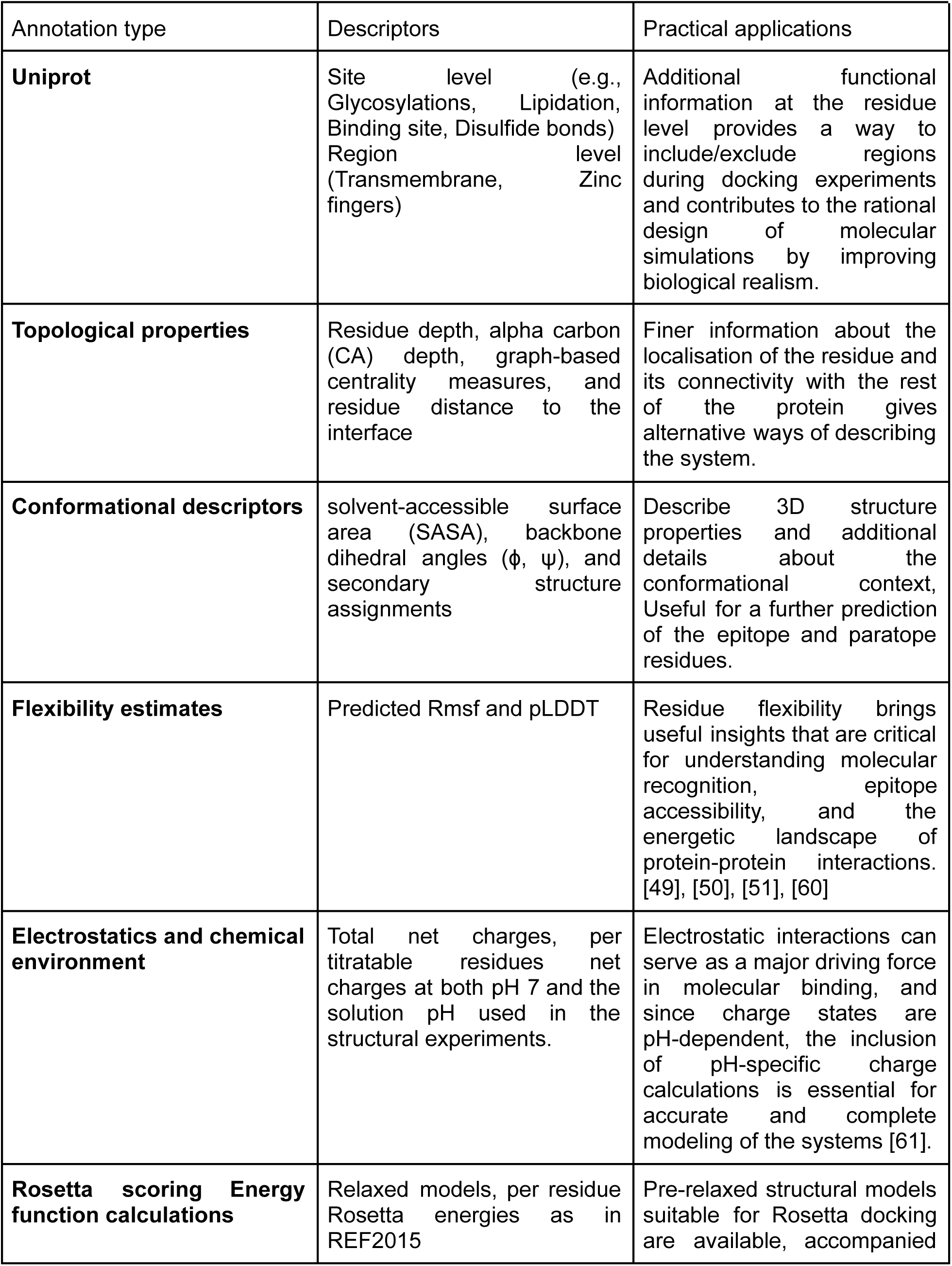
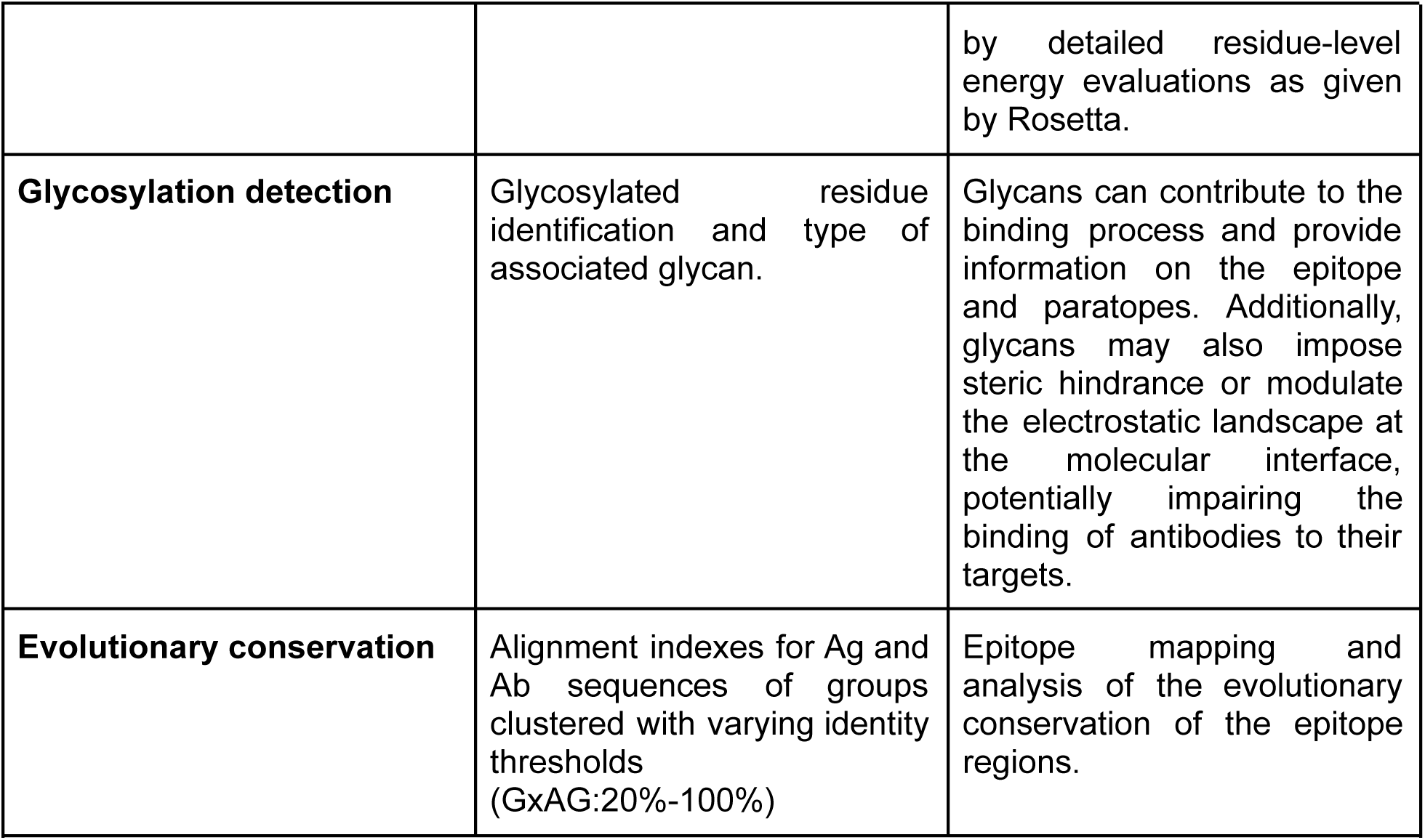
Newly computed and annotated features in the ANABAG dataset. This table categorizes annotation types, descriptors, and their respective use cases.

| Annotation type | Descriptors | Practical applications |
| --- | --- | --- |
| **Uniprot** | Site level (e.g., Glycosylations, Lipidation, Binding site, Disulfide bonds)<br>Region level (Transmembrane, Zinc fingers) | Additional functional information at the residue level provides a way to include/exclude regions during docking experiments and contributes to the rational design of molecular simulations by improving biological realism. |
| **Topological properties** | Residue depth, alpha carbon (CA) depth, graph-based centrality measures, and residue distance to the interface | Finer information about the localisation of the residue and its connectivity with the rest of the protein gives alternative ways of describing the system. |
| **Conformational descriptors** | solvent-accessible surface area (SASA), backbone dihedral angles ($\phi$, $\psi$), and secondary structure assignments | Describe 3D structure properties and additional details about the conformational context, Useful for a further prediction of the epitope and paratope residues. |
| **Flexibility estimates** | Predicted Rmsf and pLDDT | Residue flexibility brings useful insights that are critical for understanding molecular recognition, epitope accessibility, and the energetic landscape of protein-protein interactions. [49], [50], [51], [60] |
| **Electrostatics and chemical environment** | Total net charges, per titratable residues net charges at both pH 7 and the solution pH used in the structural experiments. | Electrostatic interactions can serve as a major driving force in molecular binding, and since charge states are pH-dependent, the inclusion of pH-specific charge calculations is essential for accurate and complete modeling of the systems [61]. |
| **Rosetta scoring Energy function calculations** | Relaxed models, per residue Rosetta energies as in REF2015 | Pre-relaxed structural models suitable for Rosetta docking are available, accompanied |
|  |  | by detailed residue-level energy evaluations as given by Rosetta. |
| **Glycosylation detection** | Glycosylated residue identification and type of associated glycan. | Glycans can contribute to the binding process and provide information on the epitope and paratopes. Additionally, glycans may also impose steric hindrance or modulate the electrostatic landscape at the molecular interface, potentially impairing the binding of antibodies to their targets. |
| **Evolutionary conservation** | Alignment indexes for Ag and Ab sequences of groups clustered with varying identity thresholds (GxAG:20%-100%) | Epitope mapping and analysis of the evolutionary conservation of the epitope regions. |

Our main goal was to provide an exhaustive set of features that we consider or suspect to be important to understand and further predict Ab-Ag interface characteristics. Some of the calculated features were previously considered in other datasets [35], [46]. This is the case for the **c*onformational descriptors*, i.e.,** solvent-accessible surface area (SASA), backbone dihedral angles (ϕ, ψ), and secondary structure assignments [40], [47], that have been exploited in previous studies and the **Topological properties, i.e.,** residue depth, alpha carbon (CA) depth, graph-based centrality measures, and residue distance to the interface [48]. For some of them, a distinction was made between the bound and unbound state, *i.e* for the complex and for the separated proteins extracted from the complex.

Beside these well-known characteristics, we have introduced new features like the: **Flexibility estimates:** per-residue flexibility and pLDDT predictions from Pegasus [Vander Meersche, Y., Duval, G., Cretin, G., Gheeraert, A., Gelly, J.-C., & Galochkina, T,. PEGASUS: Prediction of MD-derived protein flexibility from sequence. Protein Science, (accepted for publication)]. Indeed, the capacity of a residue to adopt different conformations has been shown to be important in binding with a partner [49], [50], [51].

##### Electrostatic properties responsive to the chemical environment

As previously mentioned, pH and salt concentration can play a critical role in the binding process [52]. To account for these factors, we computed per-residue and total net charges of the biomolecules under physiological conditions (pH 7, 150 mM NaCl), as well as total net charges at the crystallization pH. These data were further complemented by calculations performed at physiological conditions for both bound and unbound forms of each protein [53]. Such electrostatic information is essential when assessing the performance of predictive tools.

##### Scoring of the complexes based on energy calculations

We have observed that atomic clashes may exist in some PDB structures. As a result, these structures can’t be used directly in frameworks like Rosetta, as they will have a very repulsive energy contribution, specifically due to the Lennard-Jones repulsive terms. So, we performed energy-minimization to remove these clashes according to the REF2015 Rosetta scoring energy function [54]. We provided the relaxed structures in the PDB format for download and per-residue energy scores in Rosetta energy units, derived from Rosetta’s energy minimization protocols as well as unrelaxed residue energies [55] (See the Methods section and the SM).

##### Glycosylation detection

The importance of glycosylation in protein-protein interaction is well recognized [56]. In the case of Ab or Ag, it was recently reviewed for instance by Damelang et al. [57] or Newby et al. [58]. Based on this observation, we chose to specify the glycosylation state of the residue in the dataset. We identified glycosylated residues based on the PDB structures with the Privateer software [59].

We summarized in Table 1, the types of features that were considered with their interest for analyses and prediction purposes. In addition to the annotated dataset, we provide multiple file format options and a set of modular scripts for feature extraction and analysis. More details on the choices of parameters for the tools used to calculate the feature and on how to use the information can be found in the SM.

#### Analysis of the sequence diversity of the dataset

##### Antigen sequence diversity

By clustering the complexes based on 100%, 95%, 80%, 60%, 40%, and 20% sequence identity, we created five different Ag groupings ( G100AG, G95AG, G80AG, G60AG, G40AG, G20AG). The number of sequence clusters ranges from 2,486 for G100AG to 528 for G20AG, indicating a high redundancy of Ag sequences in the starting dataset (see Table 2). The same is true for Ab sequences when considering identity thresholds below 95%.

**Table 2:** Redundancy of antigen sequences in the ANABAG dataset. Antigen sequences are grouped into clusters by a grouping method denoted as GxAG, where x represents the sequence identity threshold used for clustering (e.g., G60AG corresponds to grouping at 60% identity [x = 60]).

| Antigen Grouping (GxAG) | Number of Clusters | Mean number of sequences per cluster | Standard deviation of the number of sequences per cluster | Maximum number of sequences | Clusters with only one sequence |
| --- | --- | --- | --- | --- | --- |
| **G100AG** | 2,486 | 2.6 | 10.0 | 381 | 1,506 |
| **G95AG** | 1,255 | 5.1 | 42.2 | 1,238 | 588 |
| **G80AG** | 958 | 6.7 | 49.8 | 1,262 | 391 |
| **G60AG** | 832 | 7.7 | 57.8 | 1,288 | 321 |
| **G40AG** | 719 | 8.9 | 63.8 | 1,288 | 262 |
| **G20AG** | 528 | 12.2 | 79.1 | 1,346 | 176 |

**Table 3:** Redundancy of antibody sequences in the ANABAG dataset. Antibody sequences are grouped into clusters by a grouping method denoted as GxAB, where x represents the sequence identity threshold used for clustering (e.g., G95AB corresponds to grouping at 95% identity [x=95]).

| Antigen Grouping (GxAB) | Number of Clusters | Mean number of sequences per cluster | Standard deviation of the number of sequences per cluster | Maximum number of sequences | Clusters with only one sequence |
| --- | --- | --- | --- | --- | --- |
| **G100AB** | 3,913 | 1.6 | 7.6 | 340 | 3,105 |
| **G95AB** | 2,780 | 2.3 | 11.5 | 497 | 1,873 |
| **G80AB** | 750 | 8.6 | 156.6 | 4,279 | 459 |
| **G60AB** | 15 | 430.1 | 1,658.7 | 6,426 | 8 |

The large standard deviation of the number of sequences within clusters in several groupings illustrates the existence of unique complexes as well as high redundancy. For instance, some groups are composed of hundreds of sequences, like for the Spike 2 protein of SARS-CoV-2, which has 381 representatives in G100AG grouping, whereas the dataset contains 528 groups of sequences sharing less than 20% sequence identity (see Table 2). Among them, 176 groups are composed of only one Ag sequence. This highlights the relative diversity of antigens in the dataset, as well as the contrast between extensively studied targets and those that are either less characterized or more challenging to resolve experimentally.

##### The interface sequence diversity

The sequence identity evaluated separately for each component of the complex in no way prejudges the diversity of the interface. We thus examined the diversity of composition of interfaces by calculating the epitope index overlap (CIO) and the similarity of epitope-paratope amino acid frequencies (Sc) between pairs of interfaces. This method allows for the distinction of complexes based on the epitope region and the interface amino acid composition.

Considering different percent identity thresholds for clustering antigen sequences (G100AG to G20AG), we evaluate the number of non-redundant/unique interfaces in each cluster (see Table 4). For each cluster, the alignment index of each sequence and the interface composition of each corresponding complex were extracted (as a vector of 40 amino acid frequencies, 20 for Ab, 20 for Ag, see the details in the Methods section). Within each cluster, all pairs of interfaces were compared and grouped based on CIO and Sc. The number of resulting groups from an Ag sequence cluster gives the number of unique interfaces in this cluster. As shown in Table 4, the diversity of interfaces consistently exceeds the diversity of Ag sequences across all GxAG grouping methods. This suggests that the dataset contains multiple distinct paratope–epitope associations for the same Ag.

**Table 4:** Number of unique interfaces per antigen group, based on the Coefficient of Interface Overlap (CIO) and cosine similarity (Sc). The table relates the number of distinct interfaces to the number of sequence-based clusters, illustrating the added value of incorporating both sequence and interface redundancy in the dataset.

| Antigen Grouping (GxAG) | Number of Clusters | Number of distinct interfaces | Mean number of distinct interfaces per cluster | Standard deviation of distinct interfaces per cluster | Clusters with only one interface |
| --- | --- | --- | --- | --- | --- |
| **G100AG** | 2,486 | 4,641 | 1.87 | 5.89 | 1,847 |
| **G95AG** | 1,255 | 4,224 | 3.37 | 18.80 | 710 |
| **G80AG** | 958 | 4,219 | 4.40 | 23.26 | 482 |
| **G60AG** | 832 | 4,206 | 5.06 | 29.79 | 406 |
| **G40AG** | 719 | 4,206 | 5.85 | 34.01 | 331 |
| **G20AG** | 528 | 4,206 | 7.96 | 42.52 | 215 |

The number of unique interfaces decreases with the percentage identity threshold used for clustering, as seen in Table 4. This is expected as some complexes have a very similar interface in terms of epitope localisation and amino acid composition, while their associated Ag sequences are defined as dissimilar, given their sequence identity and the identity threshold used for grouping. These results highlight that distinct antigenic sequences can have identical interfaces in terms of epitope sequence region and amino acid composition. Conversely, by applying a redundancy criterion based on the interface similarity and the sequence identity, we can include complexes with similar Ag or Ab sequences that have distinct interfaces.

#### Localizing missing residues with respect to the interface

Among the 6,452 complexes, Ag structures were missing more residues than Ab structures. The analysis shows that a large number of gaps are located far from the interface (73% at more than 25Å), which means that the modelled residues would not have a major impact on the further analyses of the interface. Nevertheless, we observed that 5% of the missing regions are located within 10 Å of the interface, with less than 2% corresponding to gaps of 10 residues. Among the regions situated within this 10 Å threshold, 28.4% (1.4% of the total) contain more than 10 missing residues, 18.6% have between 5 and 10, 35.7% between 3 and 5, and 17.3% consist of 1 or 2 missing residues. We proceeded to complete these gaps using Modeller v10.6 [37],and the distribution of completed gaps relative to their distance to the interface is shown in Figure S1.

This analysis reveals that missing regions of large size can occur in close proximity to the binding interface. These absent residues likely correspond, in many cases, to flexible or disordered segments. While the decision to model such regions may be subject to debate, it is important to note that their occurrence is limited within our dataset. More critically, several computed properties depend on residue-level information regardless of precise atomic detail. Their omission might have a large impact on the calculated or predicted properties. This is particularly true for pH-dependent properties, such as net protein charge. Fortunately, as these calculations are based on a coarse-grained titration model here, they are less affected by local conformational variability. This observation further supports the inclusion and need of modeled residues to improve the accuracy of these theoretical estimations.

### Antigen Glycosylation

Glycan analysis presents a complex challenge due to the structural and binding site diversity of glycosylation. Glycans differ not only in their saccharide composition and stereochemistry, but also in the specific sites at which they bind to proteins. Even for a single antigenic binding site, the glycosylation pattern can vary depending on the types of enzymes responsible for the post-translational modification. It is itself influenced by the cellular compartment and the expression system used to produce the protein [56], [62]. Additionally, large and flexible glycans can hinder protein crystallization and even lead to crystallization failure [63]. As the presence of glycosylation and their diversity at the site level depend on the many previously mentioned factors, it greatly complicates their systematic analysis. The inclusion of glycan annotations in the dataset is particularly relevant, as glycans can serve dual roles in Ab-Ag interactions. On one hand, they may contribute to the binding process by shaping or stabilizing epitope and paratope conformations. On the other hand, they can impose steric hindrance or alter the local electrostatic environment at the interface, thereby impairing Ab access or changing binding affinity. As such, having access to glycosylation data enables a more accurate modeling of interaction dynamics and improves the reliability of predictive approaches, particularly in docking and epitope mapping workflows, especially for the glycan-mediated interfaces or the heavily glycosylated proteins.

Our analyses of ANABAG demonstrate that glycans can indeed affect binding, either by a potential shielding effect, steric hindrance, or even by being directly targeted by the Ab. These findings highlight the need to consider glycosylation in both epitope prediction and Ab design. Among the 6,452 individual complexes in our dataset, 2,511 Ags and 136 Abs were glycosylated. A total of 15,344 glycosylation sites in Ags were identified at the structural level using the Privateer software [59], while 142 were found in Abs. Of the Ag glycosylations found at the structural level, 11,382 were also annotated at the UniProt sequence level, whereas 3,962 were detected at the structural level and were not annotated by UniProt. Additionally, 16,678 glycosylation annotations from UniProt were not observed at the structural level. On average, glycosylated Ags contained 6.1 glycosylation sites, whereas Abs had only 1.0. The maximum number of glycosylation sites detected on a single structure was 75 for Ags and 3 for Abs (for a summary of this data, see Table S2).

How these glycosylations could interfere with the interface is a major question, but difficult to settle, since we examined the complexes once formed. The distance between glycosylated residues and the interface is represented in Figure 3. We observed that 7% of glycosylated residues can be found at less than 10Å from an Ab residue in the interface and 34% between 10 to 25Å. Depending on the length of the sugar, it might indeed have an impact on the interface. Conversely, the rest of the glycan carrying residues are found at distances greater than 25Å, a distance that probably prohibits the glycan from directly contributing to the interface. Despite not being directly involved in the interface, they can still exert influence via significant electrostatic coupling with titratable residues in the vicinity.

**Figure 3:**
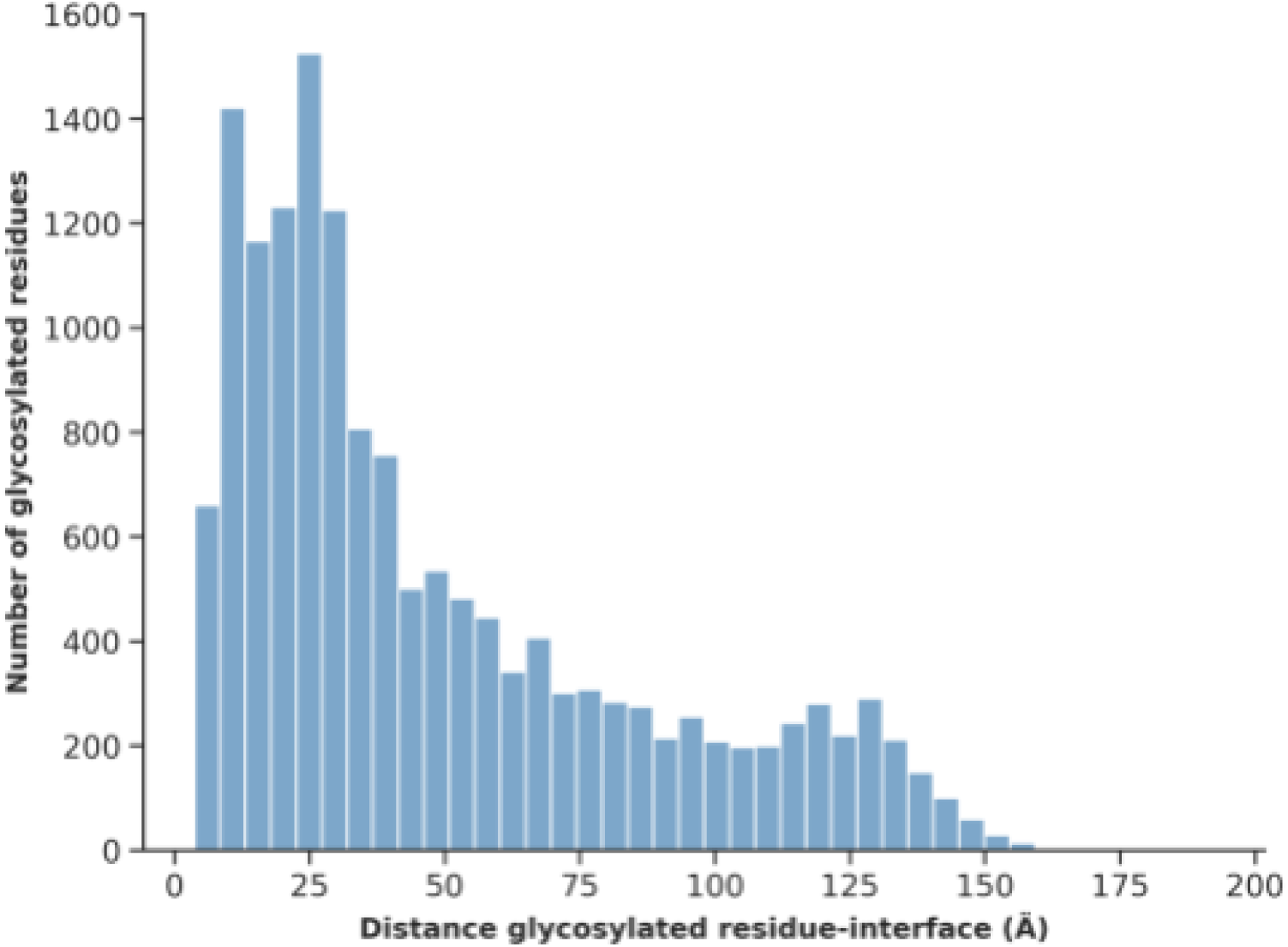
Distribution of glycosylated residues in antigens as a function of their proximity to the Ag–Ab interface. The distance is defined as the shortest spatial separation between the center of mass of each glycosylated residue and any residue located at the interface.

#### Evaluating the shielding effect

The analysis detailed above shows that Ab-Ag interfaces are generally located outside of glycosylated regions. However, certain Abs specifically target glycans. Indeed, in ANABAG, we found that over the 6,452 antibodies, 1,400 made at least one contact (distance between glycan and paratope atom < 5Å) with a glycan, and the most contacted glycan is the N-acetyl-beta-D-glucosamine (NAG) (see Table 5). Some of these interfaces are glycan-mediated, which means that the interface is primarily composed of glycan-paratope contacts. This is the case for the envelope glycoprotein gp160 of HIV presented in Figure 4A, where glycans play a dominant role in binding and make more than 75% of the total contacts with the Ab (see Figure 4A). In these cases, the glycan part of the Ag is targeted by the Ab, and such examples must be taken into consideration, especially for tasks such as epitope prediction or docking.

**Figure 4:**
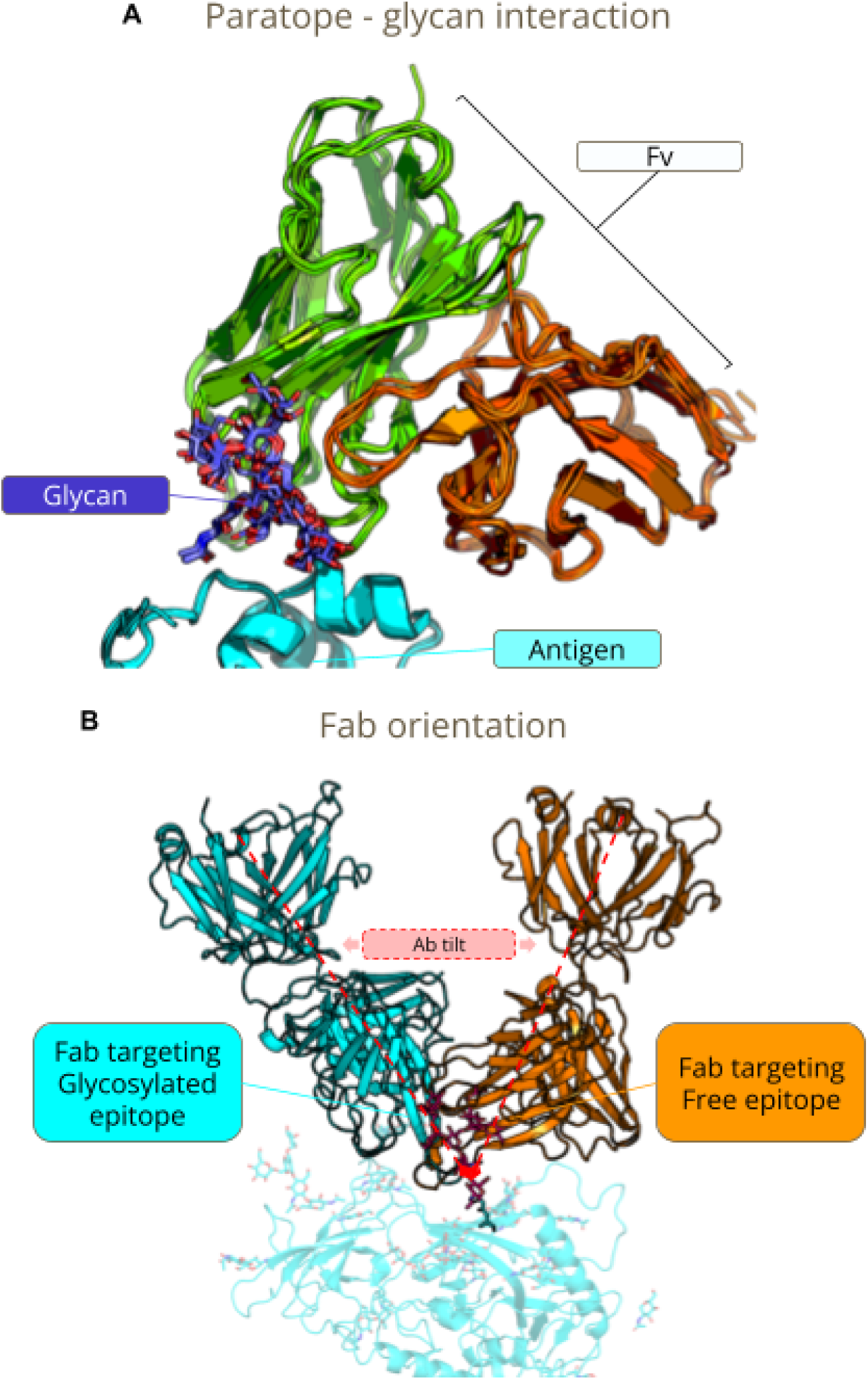
Glycan-driven modulation of Ag-Ab interactions. (A) Top panel: Structural illustration of a fragment Ab (green and orange) targeting a glycosylation site (purple) on the surface of an Ag (cyan). The depicted complexes are derived from PDB entries: 6mtj_0, 6mtn_1, 6mu6_1, 6mu7_1, 6mu8_0, 6nm6_0, 6nnf_1, 6nnj_2, 6opa_0, and 8d0y_0. (B) Bottom panel: Antibody orientation shift caused by glycan-induced steric hindrance. The red dotted cone highlights the angle between the two conformations. The glycosylated antigen is shown as a cartoon with transparent cyan glycans. The glycan responsible for the tilt is shown in hot pink sticks. The Ab bound to the glycosylated complex is in cyan; the antibody in the glycan-free complex is shown in orange. Structures are derived from PDB IDs: 3jcb_0 and 4nco_1.

**Table 5:**
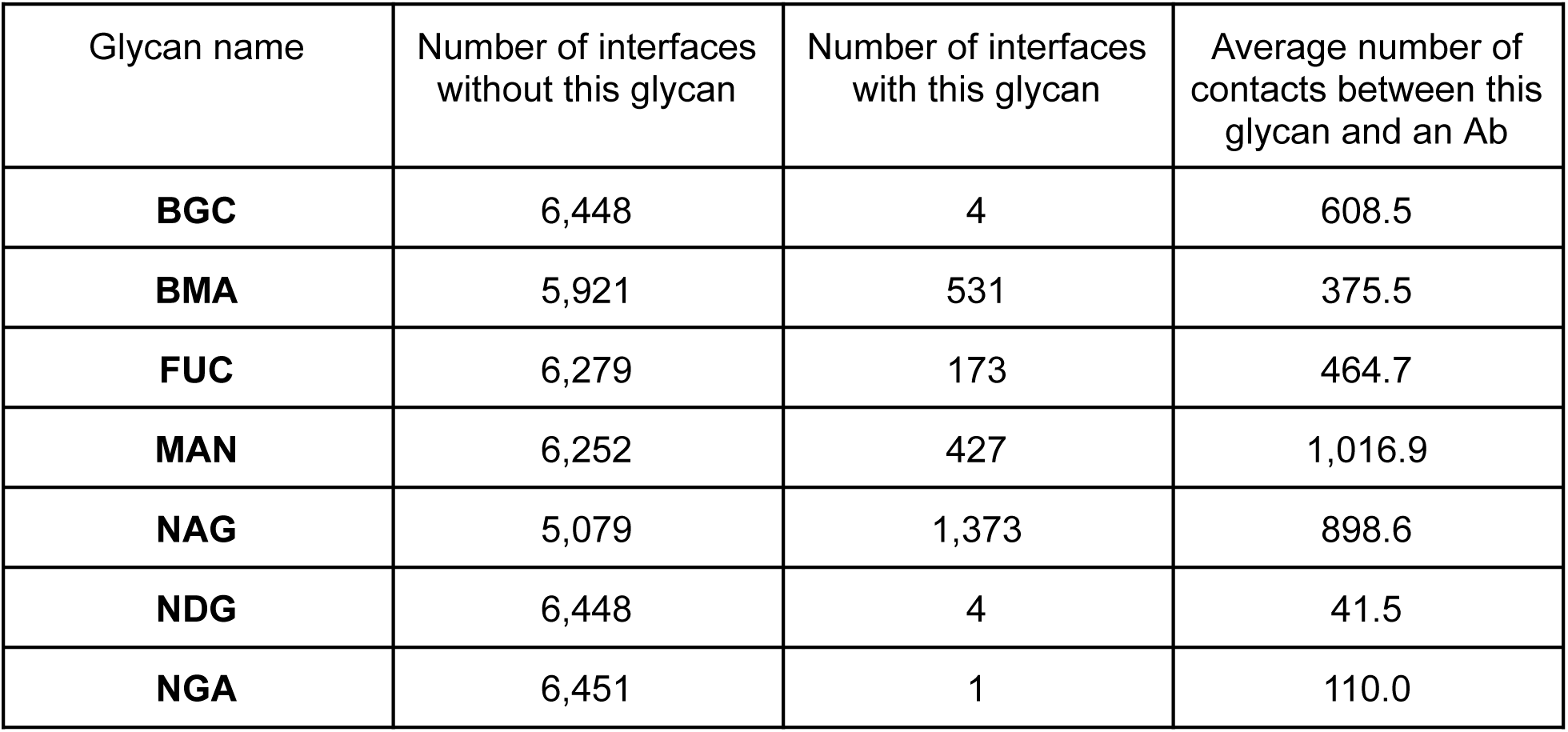
Summary of glycan-mediated contacts across the 6,452 Ag–Ab complexes in the dataset. For each glycan type, the table reports the number of interfaces in which the glycan establishes contact (interface with this glycan) and those in which it is absent (interface without this glycan), as well as the average number of heavy-atom contacts formed between Ab residues and the glycan. The average was computed by dividing the total number of Ab–glycan contacts by the number of glycan-present interfaces. Abbreviations: BGC – β-D-Glucose; BMA – β-D-Mannose; FUC – L-Fucose; MAN – α-D-Mannose; NAG, NDG – N-Acetylglucosamine; NGA – N-Acetyl-D-galactosamine.

| Glycan name | Number of interfaces without this glycan | Number of interfaces with this glycan | Average number of contacts between this glycan and an Ab |
| --- | --- | --- | --- |
| **BGC** | 6,448 | 4 | 608.5 |
| **BMA** | 5,921 | 531 | 375.5 |
| **FUC** | 6,279 | 173 | 464.7 |
| **MAN** | 6,252 | 427 | 1,016.9 |
| **NAG** | 5,079 | 1,373 | 898.6 |
| **NDG** | 6,448 | 4 | 41.5 |
| **NGA** | 6,451 | 1 | 110.0 |

In ANABAG, we have also identified examples of highly similar Ag sequences in the same G95AG group that are, in some cases, glycosylated and in others glycan-free. These complexes offer the opportunity to evaluate the effect of glycan on the conformation of the complex and the nature of the interface. A central question is whether glycosylation contributes to shielding the epitope and thereby modulates Ab-Ag binding? Recent studies, particularly on the glycosylated SARS-CoV-2 spike protein, have challenged the notion that glycan shielding completely prevents Ab binding due to their additional roles and flexibility [64], [65], [66]. Indeed, both Abs and glycans are flexible molecules, allowing them to accommodate steric hindrances through small structural adjustments. We confirmed this adaptability by analyzing two Ag-Ab complexes with identical sequence (100% sequence identity); one with a glycosylated epitope and the other without.

Figure 4B shows that the glycan presence introduces a tilt in the Ab orientation, also modifying the epitope composition. This is highlighted by our interface diversity measure, which labels these two interfaces as different in terms of amino acid composition. However, overall, the location of the binding region on the surface of the protein remains roughly unchanged, and the presence of the glycan is the only structural component that accounts for the change in the Ab orientation. This is a clear example of two complexes with identical Ag sequences that exhibit different binding conformations due to the presence of a glycan found using the ANABAG dataset. In this case, the Ab is not shielded or redirected to a distant epitope; instead, it targets the same region but adopts a constrained binding orientation as a result of the glycan’s steric hindrance. This observation reinforces the idea that paratopes are highly specific to a given epitope region and can still bind to them, regardless of glycosylation.

### Examples of paratopes with different amino acid compositions binding the same epitope

We examined to what extent the data available in ANABAG informs about the diversity of Ab sequences targeting the same epitope. We identified 78 epitopes that are each targeted by two or more Abs sharing less than 95% sequence identity. For each epitope, we assessed the similarity among the paratopes that recognized it by calculating the cosine similarity between their amino acid frequencies (see Figure 5A).

**Figure 5:**
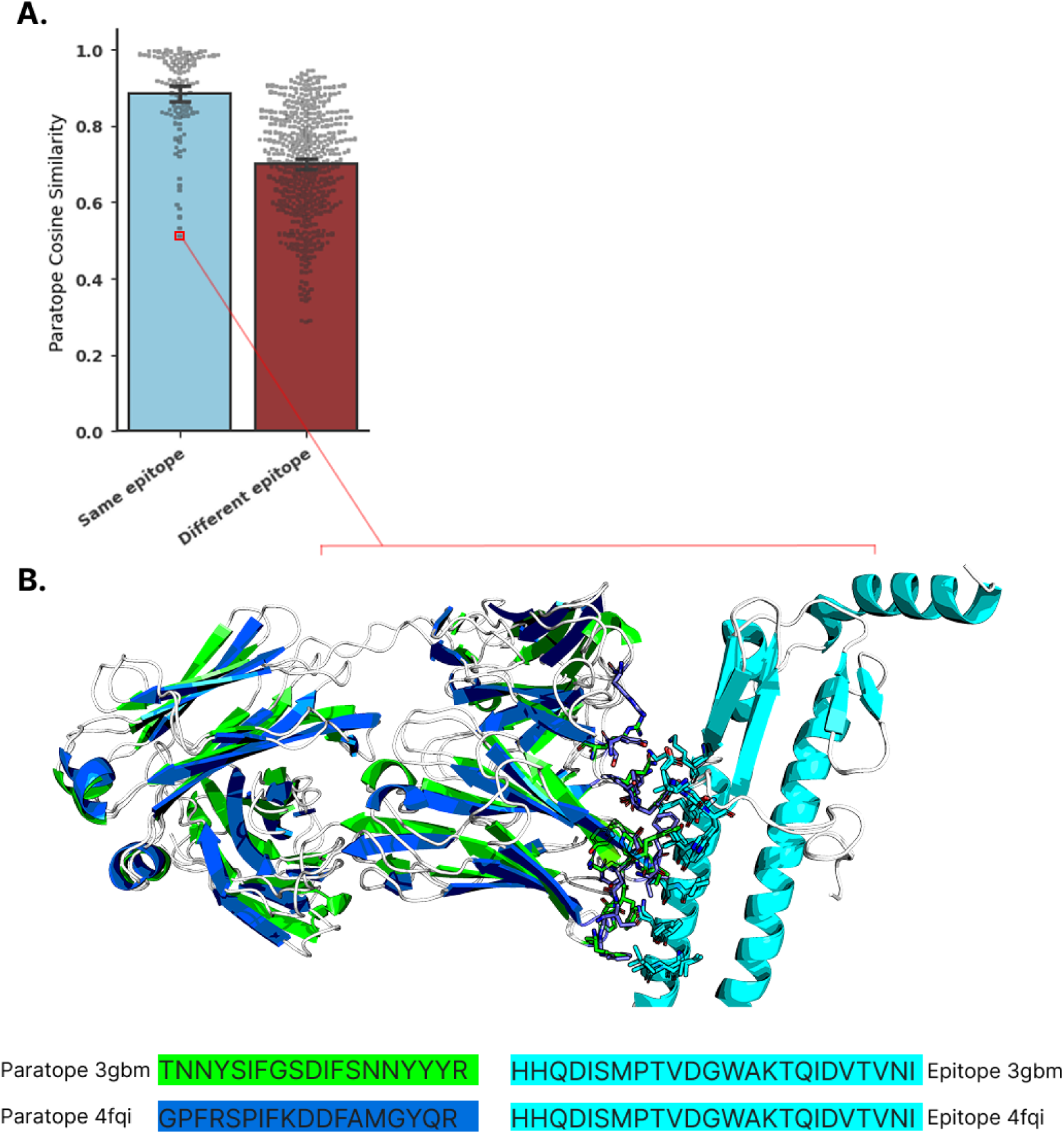
Diversity of paratopes that recognize the same epitope. (A) Top panel: Distribution of cosine similarity (CS) values for amino acid composition in paratope pairs, distinguishing those that recognize the same versus different Ags (based on G100AG clustering). The red square marks the example highlighted in panel B. (B) Bottom panel: Structural illustration of two Fab fragments (PDB IDs: 3gbm_0 and 4fqi_0) that share less than 95% sequence identity and display distinct paratope compositions, yet both recognize the same epitope region. The compositions of Paratopes are highlighted in green and blue, and the epitopes in cyan. Paratopes and epitopes alignments are included in the SM.

For these specific pairs of paratopes that share less than 95% identity and recognize the same epitope, their amino acid composition is highly similar, as shown in Figure 5A. Indeed, the average amino acid similarity for paratopes targeting the same antigen is 0.9. As a baseline for comparison, the cosine similarity of Abs paratopes targeting different Ags is smaller (∼0.7), which means a higher dissimilarity (see Figure 5A).

Conversely, we also observe some pairs of paratopes with distinct amino acid composition that bind to the same epitope of the exact same Ag. We illustrate this by a specific example with the hemagglutinin of influenza virus H5N1 that has an epitope which is recognized by two Abs sharing less than 95% sequence identity. The two paratopes recognize the same epitope despite the fact that their amino acid composition is dissimilar, with a cosine similarity of ∼0.5 (Figure 5B).

These results highlight that distinct paratope compositions can, in some cases, converge on the same epitope region. However, such cases appear to be exceptions, as when two distinct antibodies - sharing less than 95% identity - recognize the same epitope, their paratope is usually highly similar.

This broader observation suggests that most Ab–Ag interfaces in ANABAG follow a lock-and-key mechanism, in which the epitope and paratope compositions are highly specific to one another—typically interacting exclusively—despite variation in the sequence outside the interface. As seen in Figure 5A, most pairs of paratopes targeting the same epitope composition are similar despite that their corresponding Ab sequence shares less than 95% identity. This observation suggests that Ab-Ag binding properties are less affected by sequence changes outside of the binding region, which is in line with what is observed by others [12], [67]. As an important note, in the case of VHH, - single domain antibodies - the paratope and CDR should be distinguished because VHH tend to have more paratope residues outside of the CDR regions compared to the “classical” antibodies [14], [68].

### Antibody polyspecificity

In the previous section, we focused on the diversity of paratopes able to recognize a given epitope. In the present section, we focus on Ab polyspecificity, which corresponds to the capacity of an Ab to bind different targets with varying degrees of affinity. For most of the complexes, we don’t have any experimental binding affinity measures, which would give a quantitative understanding of Ab polyspecificity and would allow us to rank different complexes based on it. Still, we can assess the diversity of antigenic targets for 100% identical antibodies in ANABAG.

We identified all Abs binding to multiple Ags belonging to different sequence clusters, based on the GxAG schemes. A polyspecific Ab, in the context of a given GxAG, is defined as one that binds to at least two Ags from different clusters within that GxAG.

In Table 6, we considered the different groups built on the sequence identity of the Ags and detailed quantitative data that illustrate this polyspecificity. For instance, in the G100AG group that contains 2,486 clusters, we identified 3,574 Abs targeting one specific Ag but 485 targeting 2 distinct Ag or more. It is also found that the same antibody can target 69 different targets. Since the distinct clusters at this identity threshold could be very similar, this polyspecificity might be discussed. However, for the lowest threshold, in the G20AG (Ag sequences sharing less than 20% identity) that contains 528 distinct clusters, we still found 8 Abs that target two or more distinct Ags, and the same antibody can target 3 very distinct Ag (Table 6).

**Table 6:** Polyspecificity of Abs across different Ag grouping schemes in the ANABAG dataset. Antibody reactivity is assessed based on the number of distinct antigen clusters (GxAG) each Ab binds to, where x denotes the sequence identity threshold used for Ag clustering.

| Antigen Grouping (GxAG) | Number of antibodies targeting one specific antigen | Number of antibodies targeting two or more distinct antigens | The maximum number of distinct antigens targeted by one antibody |
| --- | --- | --- | --- |
| **G100AG** | 3,574 | 485 | 69 |
| **G95AG** | 3,910 | 149 | 6 |
| **G80AG** | 3,979 | 80 | 5 |
| **G60AG** | 4,032 | 27 | 5 |
| **G40AG** | 4,048 | 11 | 4 |
| **G20AG** | 4,051 | 8 | 3 |

Overall, these results suggest that identical Abs are predominantly generated in response to the same Ag target. However, we also find evidence of Abs that bind to very different targets.

The Ag from PDB ID 7jti is shown in blue, and the one from 2vir in green. The compositions of Paratopes are highlighted in cyan, and the epitopes in green and blue. Paratopes and epitopes alignments are included in the SM.

Polyspecific antibodies target very different Ags, but this information does not provide any indication on the composition of the paratope nor the targeted epitope. Indeed, the targeted epitope could be highly similar even for different Ags. For the polyspecific Abs identified in Table 6, we compared the amino acid composition similarity between paratope and epitope pairs for the two distinct complexes in Figure 6A (two paratopes or epitopes identified from two distinct complexes where the two antibodies are identical and the two Ags are distinct).

**Figure 6:**
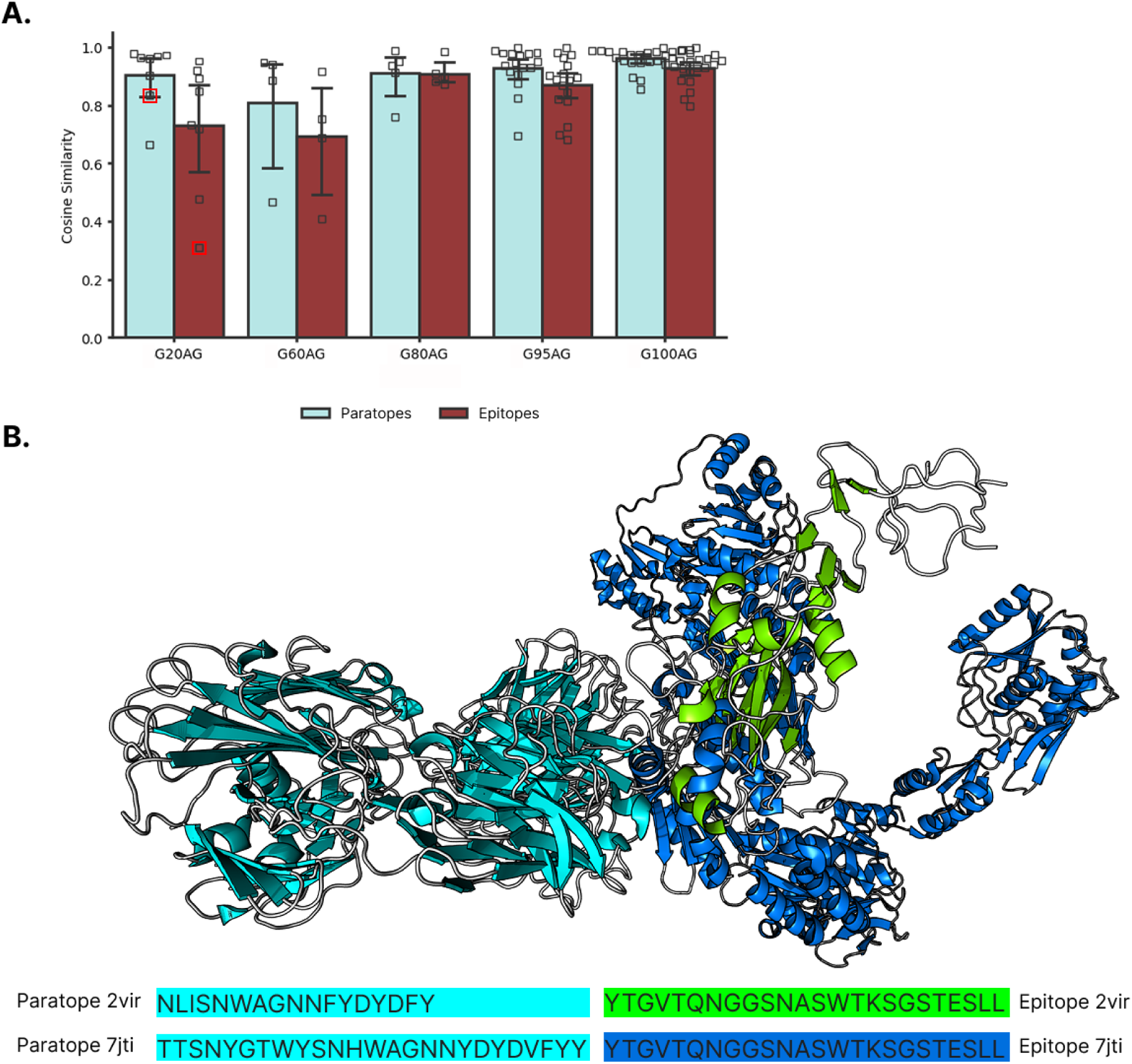
Polyspecificity of antibodies in the ANABAG dataset. (A) Top panel: Distribution of cosine similarity values for amino acid composition in interface pairs recognized by the same Ab. Red squares indicate the specific case illustrated in panel B. (B) Bottom panel: Structural example of a polyspecific Ab targeting two distinct Ags with less than 20% sequence identity.

We find that paratopes of polyspecific Abs tend to be highly similar, regardless of the Ags it targets, with an average cosine similarity of 0.9 across paratope pairs (see Figure 6A). This suggests that certain paratope compositions are broadly compatible with different Ags. In some cases, however, we observe paratope pairs from the same Ab with lower amino acid similarity, implying that antibodies may use distinct CDR combinations to bind different targets.

As an illustration, we show in Figure 6B, the structure of a Fab interacting with two dissimilar epitopes (cosine similarity of 0.30) of the Interphotoreceptor retinoid-binding protein and the hemagglutinin from the influenza virus with a highly similar paratope (cosine similarity of 0.83), amino acid composition.

When examining the epitopes recognized by these polyspecific Abs, we find that epitope pairs are, on average, more dissimilar than paratope pairs. This supports the idea that a single paratope composition can accommodate a range of epitope compositions. Nonetheless, for all Ag cluster definitions (GxAG), most epitope pairs recognized by the same Ab still exhibit high similarity, typically with a cosine similarity above 0.9 (Figure 6A).

This suggests that, despite sequence variability among Ags, polyspecific Abs in the ANABAG dataset frequently recognize epitope regions with similar compositional features.

Overall, our results show that polyspecific Abs in ANABAG largely recognize conserved epitope features across diverse Ags, although some demonstrate the capacity to adapt and bind distinct epitope compositions.

## Discussion

This study constitutes a component of a more extensive framework centred on the prediction of Ag-Ab complexes. When the three-dimensional structures of both partners are known, docking tools remain the primary approach for predicting complex formation, providing insights into binding poses and interface interactions. However, factors such as conformational flexibility, induced fit, and pH-dependent effects can affect docking accuracy [69]. In cases where structural data is unavailable, deep learning tools like AlphaFold [38] can be used to predict protein structures and complexes. They can then serve as input for docking or other computational approaches, such as molecular dynamics simulations and electrostatic analyses, to refine and evaluate interaction predictions. However, the docking procedure can be extremely time-consuming, when carried out blindly, and the success rate is rather low. Hence, any information that can help in guiding the docking is of major interest. A useful information to guide the docking process is to identify the most probable epitope residues on the Ag surface.

Indeed, accurate epitope prediction [25], [26] holds promise for various immunological applications. Instead of beginning with a specific Ab-Ag pair, this approach starts solely with the Ag characteristics, sequential or structural. Predicting which regions on the Ag surface are most likely to be recognized by Abs unlocks more efficient engineering of Abs directly targeted towards these epitopes. However, epitope prediction is a very challenging process, as some Ags have essentially their entire surface targeted by Abs, suggesting that Abs can adapt to bind to any target. Recently, methods developed by several authors including Zhu, Clifford, Tubiana, Choi and their teams [21], [22], [70], [71] have shown improvements on epitope prediction by using different predictive model architectures and features, but the success remains low. One of the difficulties in predicting epitopes originates from the extremely unbalanced number of interface residues relative to the non interface residues. Interestingly, the interface residues represent only ∼6% of the total number of Ag residues (i.e., 2,774,593 residues) in ANABAG.

So, whatever the complexity of the algorithm, the building of training datasets with informative features is the key initial step. Hence, we discuss in the next sections, the interest of the ANABAG dataset and the usefulness of important features that are not considered generally in previously built datasets.

### A dataset with original and unique features

Sabdab [29] is the landmark database in the field of structural Ab-Ag interaction. The complexes included are pre-selected and labelled with different per-complex annotation as the expression system, which protein chains are the Ab and Ag, and which ensemble of chains constitutes a separate biological unit, as well as other useful features like the experimental resolution and in some cases, the binding affinity. The structures included are not standardized and contain identical biological units in the same entry, or packing structures. However the ensemble represents a very complete view of the available Ab-Ag structural data.

From SabDab, a large number of datasets has been built with different aims, e.g. training, benchmarking purposes [9], [22], [46], [72]. The first step has generally consisted of eliminating the packing structures and standardizing the data before looking at specific cases (if needed) and calculating various features for analysis or predictive purposes. These features are usually calculated at the residue level, which SabDab lacks.

Upon building such datasets, different criteria are applied to select Ab-Ag complexes and resulting in a subset of examples that is often greatly underestimated in terms of diversity of complexes. During the construction of such datasets, a lot of different structural and sequential features are computed and available for the user. For example, Almeida et al [35] created a dataset that is a collection of structures, like SabDab, with some additional features and docked models for benchmark purposes. Kilambi et al [73] created a dataset of cognate and non cognate Ab-Ag interactions in a cross-docking experiment to evaluate their ranking according to Rosetta scores. Miller et al [46] calculated a diverse set of features and evaluated their relevance for Ab-Ag binding affinity prediction. Among the features tested, the Rosetta scoring energy function was shown to be useful in predicting the binding affinity. For this reason, we included the calculation of the Rosetta score for all systems in the present dataset, providing detailed information on its various components. In addition, we have included the *per residue* Rosetta REF2015 energies calculated from the original structure, but also from the relaxed structure in bound and unbound forms. These calculations would be useful for predicting Ab-Ag properties like binding affinity, as well as providing ready to use complexes for other Rosetta protocols (e.g., Docking) that require a minimized structure.

Among the energy components, the electrostatic interaction plays an important role, since it is a long-range term. Its contribution can even be one of the main driving forces of Ab-Ag binding [74], [75]. This is due to the higher propensity of titratable residues at the interface on the Ab and Ag side compared to non immune protein-protein interaction [23], [76], [77].

In this context, an important factor, which is frequently neglected, is the pH. The solution pH regulates the charge of the titratable residues, and at the end, all electrostatic properties and the biomolecular interactions. This factor has been used in some cases to facilitate crystallisation, favoring the stable organization of the molecules in the crystal cell. For instance, in the present dataset, 17% of structures have been solved with a pH < 6.5, 8% with a pH > 8.0, which means that 25% of the structures available in the dataset have been obtained with a pH different from the physiological conditions. Understanding how this factor influences the electrostatic properties of both partners is crucial before generalizing any trends. To accurately represent the electrostatic characteristics of residues and structures in ANABAG, we chose to calculate the charges of all titratable aminoacids using the FPTS [53], considering the solution pH, salt concentration and the local environmental influence. The method is a titration scheme based on Monte Carlo simulations [78] that enables fast and accurate prediction of the electrostatic properties of biomolecules. It has demonstrated success in benchmark studies [79] and has already been applied to characterize viral epitopes [80]. Here, the charges, dipole moments and the charge regulation capacity [81] have been calculated at the experimental pH and at pH 7.0 for the Ab-Ag complexes and for the Ab and Ag separately, extracted from the complexes. A rapid analysis shows that, as expected, in most cases when the experimental pH is close to the physiological pH, there are no major changes in the charges of the titratable residues. However, when the experimental pH differs by at least two pH units from the physiological pH, the impact is significant. For example, the Major Pollen Allergen bet v 1-a, in complex with an antibody (PDB id 1FSK_2), contains a large number of titratable residues. The X-ray structure was solved at pH 4.0. This results in differences in charge of more than 0.6 charge units for ASP and GLU residues. The total charge of each chain is therefore significantly different to what it would be at pH 7.0. This may affect the way the two partners interact. Also, for interface properties, the net charge of the interfaces can be misrepresented by more than 4 units if the calculation is made at pH 7.0 instead of the experimental pH. This becomes critical for complexes where the electrostatic interaction is the main driving force, and the inclusion of a pH-dependent charge is of great value to accurately represent these types of Ab-Ag interactions.

Consequently, these charges that are given in the dataset constitute additional features that can be valuable for a better understanding of the Ab-Ag association and useful starting points for further simulations at the experimental structural pH and at pH 7.0.

### Useful tools to quantify interface similarity

Besides the *per residue f*eatures, we also propose groupings of complexes based on the interface similarity, quantified with some measures, the CIO & Sc that can be useful for building new datasets. This feature is of interest to take into account the epitope immunodominance [82], [83], i.e., an epitope region that can be recognized by different Abs and polyspecificity, which are often overlooked when creating datasets. Hence, the information will help to identify if the targeting paratopes share similarity. The process of building docking datasets could benefit greatly by using interface similarity as a criterion during the structure selection process. This way, both polyspecific and immunodominant epitope examples could be included, and these cases are critical for understanding epitope-paratope pairings and distinguishing binders from non-binders.

Despite their potential involvement in Ab binding, glycans are largely overlooked in most Ab-Ag analyses and prediction methods, as stated by Cia et al [26]. Many computational tools for epitope prediction, (such as ScanNet [22], and BepiPred [21]) include epitope information derived from glycan-mediated complexes, such as the HIV envelope glycoprotein gp160, without explicitly considering the presence of the glycan’s atoms. These cases can lead to misleading predictions, as upon glycan removal of glycan-mediated interfaces, the interface may remain largely empty or incorrectly defined.

Another difficulty stems from the possibility of an Ag to carry multiple epitopes, making the classification of epitope residue more tricky. A strategy employed by Clifford et al [21] called epitope collapse, consists of mapping all epitope residues of an ensemble of sequences in an MSA onto a representative sequence. However, this method is sensitive to the sequence identity threshold that is selected for defining what sequences are similar, and doesn’t take into account the variability of sequences, structure, and experimental conditions for these mapped epitopes.

With ANABAG, the user can apply a similar epitope collapse strategy by analysing the differences in all provided features between the source epitope residue and the mapped sequence, across different sequence identity thresholds. This will provide a much more sensitive evaluation of residue similarity between sequences and a more robust epitope residue identification.

## Conclusions

In this work, we introduce ANABAG, a rigorously curated collection of 6,452 Ab-Ag complexes derived from SabDab. Each structure has undergone thorough preprocessing: gaps of up to 20 residues were modeled with Modeller; UniProt annotations (glycosylation sites, transmembrane segments, zinc-finger motifs, etc.) were mapped; and numerous unique to this dataset residue-level features like glycosylations and pH dependent physical chemical properties (net charge, dipole moment and charge regulation capacity) were computed. Unlike many existing datasets, ANABAG preserves all unique biological units within each crystal, ensuring that structurally or functionally distinct Ab-Ag pairings (in the same PDB entry) are retained for downstream selection and analysis. Users can filter by any combination of Ag identity, Ab identity, interface redundancy, glycosylation status, or any other included feature, enabling easy and modular dataset assembly for diverse tasks from epitope prediction to benchmarking docking algorithms. ANABAG can be interrogated with integrated scripts to select Ab-Ag complexes based on any of the calculated or annotated features, sequence identity, and interface similarity.

## Acknowledgments

This work has been supported by the “Agence Nationale de la Recherche” [ANR-20-CE06-0029] within the frame of the Emulate project “Developing, Validating and applying computer simulation methods to Enhance the MolecULAr undersTandIng and tO eNgineer functionalized biomaterials” – EMULATE and by the “Fundação de Amparo à Pesquisa do Estado de São Paulo” [Fapesp 2020/07158–2 (F.L.B.d.S.)] and the Conselho Nacional de Desenvolvimento Científico e Tecnológico (CNPq) [CNPq 305393/2020–0 (FLBdS)]. We thank the Très Grand Centre de Calcul for giving us access to their supercomputer [Projects A0140714172, AD010713871, AD010710878R1]. We would also like to thank Bilal Delikaya for his assistance in computing several features of the dataset.

## Supporting Informations

Additional scripts and support for selecting cases based on the dataset features can be downloaded at https://github.com/DSIMB/anabag-handler.git

## AUTHOR INFORMATION

Notes The authors declare no competing financial interest.

## References

[1] M. H. V. Van Regenmortel, « Specificity, polyspecificity, and heterospecificity of antibody-antigen recognition », J. Mol. Recognit., vol. 27, n° 11, p. 627-639, nov. 2014, doi: 10.1002/jmr.2394.

[2] C. Janeway, Éd., Immunobiology 5: the immune system in health and disease, 5th ed. New York: Garland Pub, 2001.

[3] Y. Elhanati, Z. Sethna, Q. Marcou, C. G. Callan, T. Mora, et A. M. Walczak, « Inferring processes underlying B-cell repertoire diversity », Philos. Trans. R. Soc. B Biol. Sci., vol. 370, n° 1676, p. 20140243, sept. 2015, doi: 10.1098/rstb.2014.0243.

[4] R. R. Porter, « The hydrolysis of rabbit γ-globulin and antibodies with crystalline papain », Biochem. J., vol. 73, n° 1, p. 119-127, sept. 1959, doi: 10.1042/bj0730119.

[5] G. M. Edelman et M. D. Poulik, « Studies on structural units of the gamma-globulins », J. Exp. Med., vol. 113, n° 5, p. 861-884, mai 1961, doi: 10.1084/jem.113.5.861.

[6] G. M. Edelman, B. A. Cunningham, W. E. Gall, P. D. Gottlieb, U. Rutishauser, et M. J. Waxdal, « THE COVALENT STRUCTURE OF AN ENTIRE γG IMMUNOGLOBULIN MOLECULE », Proc. Natl. Acad. Sci., vol. 63, n° 1, p. 78-85, mai 1969, doi: 10.1073/pnas.63.1.78.

[7] T. T. Wu et E. A. Kabat, « AN ANALYSIS OF THE SEQUENCES OF THE VARIABLE REGIONS OF BENCE JONES PROTEINS AND MYELOMA LIGHT CHAINS AND THEIR IMPLICATIONS FOR ANTIBODY COMPLEMENTARITY », J. Exp. Med., vol. 132, n° 2, p. 211-250, août 1970, doi: 10.1084/jem.132.2.211.

[8] P. Bork, L. Holm, et C. Sander, « The Immunoglobulin Fold », J. Mol. Biol., vol. 242, n° 4, p. 309-320, sept. 1994, doi: 10.1006/jmbi.1994.1582.

[9] P. B. P. S. Reis, G. P. Barletta, L. Gagliardi, S. Fortuna, M. A. Soler, et W. Rocchia, « Antibody-Antigen Binding Interface Analysis in the Big Data Era », Front. Mol. Biosci., vol. 9, p. 945808, juill. 2022, doi: 10.3389/fmolb.2022.945808.

[10] K. Masuda, K. Sakamoto, M. Kojima, T. Aburatani, T. Ueda, et H. Ueda, « The role of interface framework residues in determining antibody V_H_ /V_L_ interaction strength and antigen-binding affinity », FEBS J., vol. 273, n° 10, p. 2184-2194, mai 2006, doi: 10.1111/j.1742-4658.2006.05232.x.

[11] W.-C. Liang et al., « Dramatic activation of an antibody by a single amino acid change in framework », Sci. Rep., vol. 11, n° 1, p. 22365, nov. 2021, doi: 10.1038/s41598-021-01530-w.

[12] G. L. Gordon, H. L. Capel, B. Guloglu, E. Richardson, R. L. Stafford, et C. M. Deane, « A comparison of the binding sites of antibodies and single-domain antibodies », Front. Immunol., vol. 14, p. 1231623, juill. 2023, doi: 10.3389/fimmu.2023.1231623.

[13] E. R. Rhodes, J. G. Faris, B. M. Petersen, et K. G. Sprenger, « Common framework mutations impact antibody interfacial dynamics and flexibility », Front. Immunol., vol. 14, p. 1120582, févr. 2023, doi: 10.3389/fimmu.2023.1120582.

[14] L. S. Mitchell et L. J. Colwell, « Analysis of nanobody paratopes reveals greater diversity than classical antibodies », Protein Eng. Des. Sel., vol. 31, n° 7-8, p. 267-275, juill. 2018, doi: 10.1093/protein/gzy017.

[15] S. C. Oostindie, G. A. Lazar, J. Schuurman, et P. W. H. I. Parren, « Avidity in antibody effector functions and biotherapeutic drug design », Nat. Rev. Drug Discov., vol. 21, n° 10, p. 715-735, oct. 2022, doi: 10.1038/s41573-022-00501-8.

[16] C. Chen, Z. Garcia, D. Chen, H. Liu, et P. Trelstad, « Cost and supply considerations for antibody therapeutics », mAbs, vol. 17, n° 1, p. 2451789, déc. 2025, doi: 10.1080/19420862.2025.2451789.

[17] S. M. Paul et al., « How to improve R&D productivity: the pharmaceutical industry’s grand challenge », Nat. Rev. Drug Discov., vol. 9, n° 3, p. 203-214, mars 2010, doi: 10.1038/nrd3078.

[18] N. H. Trier et G. Houen, « Antibody Cross-Reactivity in Auto-Immune Diseases », Int. J. Mol. Sci., vol. 24, n° 17, p. 13609, sept. 2023, doi: 10.3390/ijms241713609.

[19] K. Stettler et al., « Specificity, cross-reactivity, and function of antibodies elicited by Zika virus infection », Science, vol. 353, n° 6301, p. 823-826, août 2016, doi: 10.1126/science.aaf8505.

[20] R. Akbar et al., « A compact vocabulary of paratope-epitope interactions enables predictability of antibody-antigen binding », Cell Rep., vol. 34, n° 11, p. 108856, mars 2021, doi: 10.1016/j.celrep.2021.108856.

[21] J. N. Clifford, M. H. Høie, S. Deleuran, B. Peters, M. Nielsen, et P. Marcatili, « BepiPred -3.0: Improved B-cell epitope prediction using protein language models », Protein Sci., vol. 31, n° 12, p. e4497, déc. 2022, doi: 10.1002/pro.4497.

[22] J. Tubiana, D. Schneidman-Duhovny, et H. J. Wolfson, « ScanNet: an interpretable geometric deep learning model for structure-based protein binding site prediction », Nat. Methods, vol. 19, n° 6, p. 730-739, juin 2022, doi: 10.1038/s41592-022-01490-7.

[23] G. A. Dalkas, F. Teheux, J. M. Kwasigroch, et M. Rooman, « Cation–π, amino–π, π–π, and H-bond interactions stabilize antigen–antibody interfaces », Proteins Struct. Funct. Bioinforma., vol. 82, n° 9, p. 1734-1746, sept. 2014, doi: 10.1002/prot.24527.

[24] A. V. Madsen et al., « Structural trends in antibody-antigen binding interfaces: a computational analysis of 1833 experimentally determined 3D structures », Comput. Struct. Biotechnol. J., vol. 23, p. 199-211, déc. 2024, doi: 10.1016/j.csbj.2023.11.056.

[25] X. Zeng, G. Bai, C. Sun, et B. Ma, « Recent Progress in Antibody Epitope Prediction », Antibodies, vol. 12, n° 3, p. 52, août 2023, doi: 10.3390/antib12030052.

[26] G. Cia, F. Pucci, et M. Rooman, « Critical review of conformational B-cell epitope prediction methods », Brief. Bioinform., vol. 24, n° 1, p. bbac567, janv. 2023, doi: 10.1093/bib/bbac567.

[27] H. M. Berman, « The Protein Data Bank », Nucleic Acids Res., vol. 28, n° 1, p. 235-242, janv. 2000, doi: 10.1093/nar/28.1.235.

[28] S. Ferdous et A. C. R. Martin, « AbDb: antibody structure database—a database of PDB-derived antibody structures », Database, vol. 2018, janv. 2018, doi: 10.1093/database/bay040.

[29] J. Dunbar et al., « SAbDab: the structural antibody database », Nucleic Acids Res., vol. 42, n° D1, p. D1140-D1146, janv. 2014, doi: 10.1093/nar/gkt1043.

[30] J. V. Kringelum, M. Nielsen, S. B. Padkjær, et O. Lund, « Structural analysis of B-cell epitopes in antibody:protein complexes », Mol. Immunol., vol. 53, n° 1-2, p. 24-34, janv. 2013, doi: 10.1016/j.molimm.2012.06.001.

[31] L. Xin et al., « Identification of Strategic Residues at the Interface of Antigen–Antibody Interactions by In Silico Mutagenesis », Interdiscip. Sci. Comput. Life Sci., vol. 10, n° 2, p. 438-448, juin 1, doi: 10.1007/s12539-017-0242-7.

[32] R. Yin et B. G. Pierce, « Evaluation of AlphaFold antibody–antigen modeling with implications for improving predictive accuracy », Protein Sci., vol. 33, n° 1, p. e4865, janv. 2024, doi: 10.1002/pro.4865.

[33] A. Davila et al., « AbAdapt: an adaptive approach to predicting antibody–antigen complex structures from sequence », Bioinforma. Adv., vol. 2, n° 1, p. vbac015, janv. 2022, doi: 10.1093/bioadv/vbac015.

[34] K. M. McCoy, M. E. Ackerman, et G. Grigoryan, « A comparison of antibody–antigen complex sequence-to-structure prediction methods and their systematic biases », Protein Sci., vol. 33, n° 9, p. e5127, sept. 2024, doi: 10.1002/pro.5127.

[35] D. S. Almeida et al., « AbSet: A Standardized Data Set of Antibody Structures for Machine Learning Applications », J. Chem. Inf. Model., vol. 65, n° 10, p. 4767-4774, mai 2025, doi: 10.1021/acs.jcim.5c00410.

[36] F. Sievers et al., « Fast, scalable generation of high-quality protein multiple sequence alignments using Clustal Omega », Mol. Syst. Biol., vol. 7, n° 1, p. 539, janv. 2011, doi: 10.1038/msb.2011.75.

[37] B. Webb et A. Sali, « Comparative Protein Structure Modeling Using MODELLER », Curr. Protoc. Bioinforma., vol. 54, n° 1, juin 2016, doi: 10.1002/cpbi.3.

[38] J. Jumper et al., « Highly accurate protein structure prediction with AlphaFold », Nature, vol. 596, n° 7873, p. 583-589, août 2021, doi: 10.1038/s41586-021-03819-2.

[39] The UniProt Consortium et al., « UniProt: the Universal Protein Knowledgebase in 2025 », Nucleic Acids Res., vol. 53, n° D1, p. D609-D617, janv. 2025, doi: 10.1093/nar/gkae1010.

[40] S. Mitternacht, « FreeSASA: An open source C library for solvent accessible surface area calculations », F1000Research, vol. 5, p. 189, févr. 2016, doi: 10.12688/f1000research.7931.1.

[41] S. Jones et J. M. Thornton, « Principles of protein-protein interactions. », Proc Natl Acad Sci USA, 1996.

[42] E. D. Levy, « A Simple Definition of Structural Regions in Proteins and Its Use in Analyzing Interface Evolution », J. Mol. Biol., vol. 403, n° 4, p. 660-670, nov. 2010, doi: 10.1016/j.jmb.2010.09.028.

[43] P. Jaccard, « Étude comparative de la distribution florale dans une portion des Alpes et du Jura », 1901, doi: 10.5169/SEALS-266450.

[44] J. Han, M. Kamber, et J. Pei, « Getting to Know Your Data », in Data Mining, Elsevier, 2012, p. 39-82. doi: 10.1016/B978-0-12-381479-1.00002-2.

[45] T. Calinski et J. Harabasz, « A dendrite method for cluster analysis », Commun. Stat. - Theory Methods, vol. 3, n° 1, p. 1-27, 1974, doi: 10.1080/03610927408827101.

[46] N. L. Miller, T. Clark, R. Raman, et R. Sasisekharan, « Learned features of antibody-antigen binding affinity », Front. Mol. Biosci., vol. 10, p. 1112738, févr. 2023, doi: 10.3389/fmolb.2023.1112738.

[47] W. G. Touw et al., « A series of PDB-related databanks for everyday needs », Nucleic Acids Res., vol. 43, n° D1, p. D364-D368, janv. 2015, doi: 10.1093/nar/gku1028.

[48] P. J. A. Cock et al., « Biopython: freely available Python tools for computational molecular biology and bioinformatics », Bioinformatics, vol. 25, n° 11, p. 1422-1423, juin 2009, doi: 10.1093/bioinformatics/btp163.

[49] R. Jimenez, G. Salazar, K. K. Baldridge, et F. E. Romesberg, « Flexibility and molecular recognition in the immune system », Proc. Natl. Acad. Sci., vol. 100, n° 1, p. 92-97, janv. 2003, doi: 10.1073/pnas.262411399.

[50] R. J. Blackler et al., « Antigen binding by conformational selection in near-germline antibodies », J. Biol. Chem., vol. 298, n° 5, p. 101901, mai 2022, doi: 10.1016/j.jbc.2022.101901.

[51] M. L. Fernández-Quintero, J. R. Loeffler, J. Kraml, U. Kahler, A. S. Kamenik, et K. R. Liedl, « Characterizing the Diversity of the CDR-H3 Loop Conformational Ensembles in Relationship to Antibody Binding Properties », Front. Immunol., vol. 9, p. 3065, janv. 2019, doi: 10.3389/fimmu.2018.03065.

[52] A. C. Dumetz, A. M. Chockla, E. W. Kaler, et A. M. Lenhoff, « Effects of pH on protein–protein interactions and implications for protein phase behavior », Biochim. Biophys. Acta BBA - Proteins Proteomics, vol. 1784, n° 4, p. 600-610, avr. 2008, doi: 10.1016/j.bbapap.2007.12.016.

[53] A. A. R. Teixeira, M. Lund, et F. L. Barroso Da Silva, « Fast Proton Titration Scheme for Multiscale Modeling of Protein Solutions », J. Chem. Theory Comput., vol. 6, n° 10, p. 3259-3266, oct. 2010, doi: 10.1021/ct1003093.

[54] R. F. Alford et al., « The Rosetta All-Atom Energy Function for Macromolecular Modeling and Design », J. Chem. Theory Comput., vol. 13, n° 6, p. 3031-3048, juin 2017, doi: 10.1021/acs.jctc.7b00125.

[55] S. Raman et al., « Structure prediction for CASP8 with all-atom refinement using Rosetta », Proteins Struct. Funct. Bioinforma., vol. 77, n° S9, p. 89-99, janv. 2009, doi: 10.1002/prot.22540.

[56] J. H. Prestegard, « A perspective on the PDB’s impact on the field of glycobiology », J. Biol. Chem., vol. 296, p. 100556, janv. 2021, doi: 10.1016/j.jbc.2021.100556.

[57] T. Damelang et al., « Impact of structural modifications of IgG antibodies on effector functions », Front. Immunol., vol. 14, p. 1304365, janv. 2024, doi: 10.3389/fimmu.2023.1304365.

[58] M. L. Newby, J. D. Allen, et M. Crispin, « Influence of glycosylation on the immunogenicity and antigenicity of viral immunogens », Biotechnol. Adv., vol. 70, p. 108283, janv. 2024, doi: 10.1016/j.biotechadv.2023.108283.

[59] J. Agirre, J. Iglesias-Fernández, C. Rovira, G. J. Davies, K. S. Wilson, et K. D. Cowtan, « Privateer: software for the conformational validation of carbohydrate structures », Nat. Struct. Mol. Biol., vol. 22, n° 11, p. 833-834, nov. 2015, doi: 10.1038/nsmb.3115.

[60] P. Tuffery et P. Derreumaux, « Flexibility and binding affinity in protein–ligand, protein–protein and multi-component protein interactions: limitations of current computational approaches », J. R. Soc. Interface, vol. 9, n° 66, p. 20-33, janv. 2012, doi: 10.1098/rsif.2011.0584.

[61] H.-X. Zhou et X. Pang, « Electrostatic Interactions in Protein Structure, Folding, Binding, and Condensation », Chem. Rev., vol. 118, n° 4, p. 1691-1741, févr. 2018, doi: 10.1021/acs.chemrev.7b00305.

[62] D. Watterson, N. Modhiran, et P. R. Young, « The many faces of the flavivirus NS1 protein offer a multitude of options for inhibitor design », Antiviral Res., vol. 130, p. 7-18, juin 2016, doi: 10.1016/j.antiviral.2016.02.014.

[63] P. D. Kwong et al., « Probability Analysis of Variational Crystallization and Its Application to gp120, The Exterior Envelope Glycoprotein of Type 1 Human Immunodeficiency Virus (HIV-1) », J. Biol. Chem., vol. 274, n° 7, p. 4115-4123, févr. 1999, doi: 10.1074/jbc.274.7.4115.

[64] Y. K. Choi et al., « Structure, Dynamics, Receptor Binding, and Antibody Binding of the Fully Glycosylated Full-Length SARS-CoV-2 Spike Protein in a Viral Membrane », J. Chem. Theory Comput., vol. 17, n° 4, p. 2479-2487, avr. 2021, doi: 10.1021/acs.jctc.0c01144.

[65] L. Casalino et al., « Beyond Shielding: The Roles of Glycans in the SARS-CoV-2 Spike Protein », ACS Cent. Sci., vol. 6, n° 10, p. 1722-1734, oct. 2020, doi: 10.1021/acscentsci.0c01056.

[66] Y. T. Pang, A. Acharya, D. L. Lynch, A. Pavlova, et J. C. Gumbart, « SARS-CoV-2 spike opening dynamics and energetics reveal the individual roles of glycans and their collective impact », *Commun*. Biol., vol. 5, n° 1, p. 1170, nov. 2022, doi: 10.1038/s42003-022-04138-6.

[67] S. A. McConnell et A. Casadevall, « Immunoglobulin constant regions provide stabilization to the paratope and enforce epitope specificity », J. Biol. Chem., vol. 300, n° 6, p. 107397, juin 2024, doi: 10.1016/j.jbc.2024.107397.

[68] C. Liu, H. Lin, L. Cao, K. Wang, et J. Sui, « Research progress on unique paratope structure, antigen binding modes, and systematic mutagenesis strategies of single-domain antibodies », Front. Immunol., vol. 13, p. 1059771, nov. 2022, doi: 10.3389/fimmu.2022.1059771.

[69] A. Neamtu, F. Mocci, A. Laaksonen, et F. L. Barroso Da Silva, « Towards an optimal monoclonal antibody with higher binding affinity to the receptor-binding domain of SARS-CoV-2 spike proteins from different variants », Colloids Surf. B Biointerfaces, vol. 221, p. 112986, janv. 2023, doi: 10.1016/j.colsurfb.2022.112986.

[70] J. Zhu, A. Gouru, F. Wu, J. A. Berzofsky, Y. Xie, et T. Wang, « BepiTBR: T-B reciprocity enhances B cell epitope prediction », iScience, vol. 25, n° 2, p. 103764, févr. 2022, doi: 10.1016/j.isci.2022.103764.

[71] S. Choi et D. Kim, « B cell epitope prediction by capturing spatial clustering property of the epitopes using graph attention network », Sci. Rep., vol. 14, n° 1, p. 27496, nov. 2024, doi: 10.1038/s41598-024-78506-z.

[72] K. Krawczyk, X. Liu, T. Baker, J. Shi, et C. M. Deane, « Improving B-cell epitope prediction and its application to global antibody-antigen docking », Bioinformatics, vol. 30, n° 16, p. 2288-2294, août 2014, doi: 10.1093/bioinformatics/btu190.

[73] K. P. Kilambi et J. J. Gray, « Structure-based cross-docking analysis of antibody–antigen interactions », Sci. Rep., vol. 7, n° 1, p. 8145, août 2017, doi: 10.1038/s41598-017-08414-y.

[74] N. Sinha, S. Mohan, C. A. Lipschultz, et S. J. Smith-Gill, « Differences in Electrostatic Properties at Antibody–Antigen Binding Sites: Implications for Specificity and Cross-Reactivity », Biophys. J., vol. 83, n° 6, p. 2946-2968, déc. 2002, doi: 10.1016/S0006-3495(02)75302-2.

[75] H. Wei, J. Tan, B. Zhou, X. Guan, Q. Zhong, et J. Wang, « Charged Residue Implantation Improves the Affinity of a Cross-Reactive Dengue Virus Antibody », Int. J. Mol. Sci., vol. 23, n° 8, p. 4197, avr. 2022, doi: 10.3390/ijms23084197.

[76] V. Kunik et Y. Ofran, « The indistinguishability of epitopes from protein surface is explained by the distinct binding preferences of each of the six antigen-binding loops », Protein Eng. Des. Sel., vol. 26, n° 10, p. 599-609, oct. 2013, doi: 10.1093/protein/gzt027.

[77] M. Wang, D. Zhu, J. Zhu, R. Nussinov, et B. Ma, « Local and global anatomy of antibody-protein antigen recognition », J. Mol. Recognit., vol. 31, n° 5, p. e2693, mai 2018, doi: 10.1002/jmr.2693.

[78] F. L. Barroso Da Silva, « Constant-pH Simulation Methods for Biomolecular Systems », in Comprehensive Computational Chemistry, Elsevier, 2024, p. 942-963. doi: 10.1016/B978-0-12-821978-2.00090-8.

[79] F. L. Barroso Da Silva et D. MacKernan, « Benchmarking a Fast Proton Titration Scheme in Implicit Solvent for Biomolecular Simulations », J. Chem. Theory Comput., vol. 13, n° 6, p. 2915-2929, juin 2017, doi: 10.1021/acs.jctc.6b01114.

[80] S. A. Poveda-Cuevas, C. Etchebest, et F. L. Barroso Da Silva, « Insights into the ZIKV NS1 Virology from Different Strains through a Fine Analysis of Physicochemical Properties », ACS Omega, vol. 3, n° 11, p. 16212-16229, nov. 2018, doi: 10.1021/acsomega.8b02081.

[81] F. L. Barroso Da Silva, M. Lund, B. Jönsson, et T. Åkesson, « On the Complexation of Proteins and Polyelectrolytes », J. Phys. Chem. B, vol. 110, n° 9, p. 4459-4464, mars 2006, doi: 10.1021/jp054880l.

[82] N. A. Brasher, A. Adhikari, A. R. Lloyd, N. Tedla, et R. A. Bull, « Hepatitis C Virus Epitope Immunodominance and B Cell Repertoire Diversity », Viruses, vol. 13, n° 6, p. 983, mai 2021, doi: 10.3390/v13060983.

[83] H.-O. Ito, T. Nakashima, T. So, M. Hirata, et M. Inoue, « Immunodominance of conformation-dependent B-cell epitopes of protein antigens », Biochem. Biophys. Res. Commun., vol. 308, n° 4, p. 770-776, sept. 2003, doi: 10.1016/S0006-291X(03)01466-9.

