## Supplementary Material for "ANABAG: Annotated Antibody Antigen dataset with unique features for Antibody Engineering Applications"

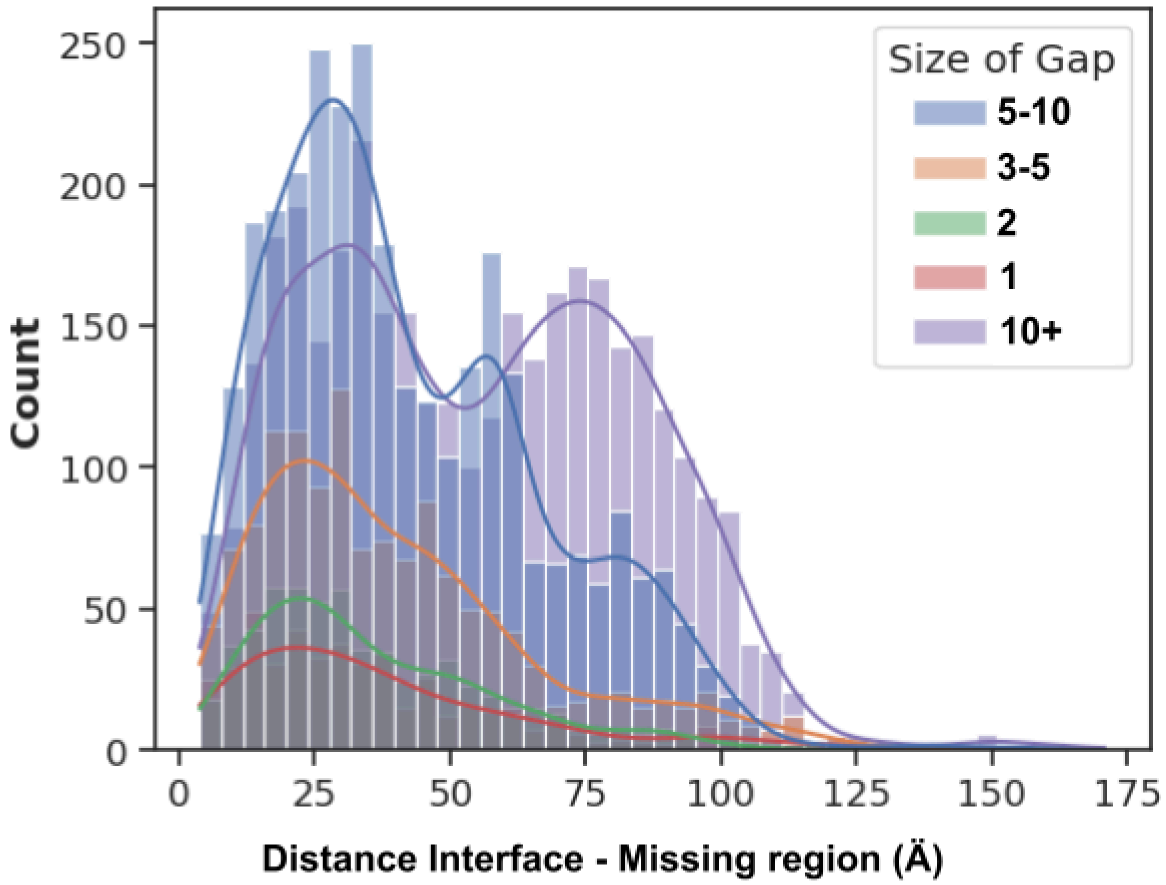


**Figure S1**: Spatial Distribution of the distance between the missing region and the Ab-Ag interface. X-axis, minimum distance between one end of the missing region and any residue center of mass in the partner. Y-axis, Number of gaps.

| Annotation type | Number of annotations | Annotations with score 5 | Annotations with score 4 | Annotations with score 3 | Annotations with score 2 | Annotations with score 1 |
|---|---|---|---|---|---|---|
| Disulfide bond | 43959 | 36839 | 2776 | 1172 | 1670 | 1502 |
| Cross link | 616 | 616 | 0 | 0 | 0 | 0 |
| Active Site | 888 | 808 | 72 | 8 | 0 | 0 |
| Binding Site | 17948 | 16937 | 574 | 182 | 184 | 71 |
| Glycosylati on | 28146 | 25115 | 875 | 365 | 429 | 1362 |
| Site | 2771 | 2680 | 85 | 4 | 2 | 0 |
| Lipidation | 314 | 301 | 13 | 0 | 0 | 0 |
| Transmem brane | 133502 | 106579 | 7469 | 2439 | 7945 | 9070 |
| Zinc finger | 856 | 782 | 74 | 0 | 0 | 0 |
| Intramemb rane | 6186 | 4492 | 1464 | 149 | 81 | 0 |

**Table S1**: Uniprot annotations for mapped antigen residues. Columns: Number of residues annotated (total and for each score). Rows: type of annotation. In each cell, the value represents the number of residues that have the corresponding row annotation for a given score (except the first column, which represents the total for all scores). While most annotations have a UniProt annotation score of 5, it is important to note that this score does not distinguish between manual and automated annotations and is not representative of the accuracy of the annotation. Rather, it represents the level of annotation of a specific entry.

| | Number of glycans detected | Total number of glycosylations detected with Privateer | Total number of glycosylations detected with Privateer and not present in Uniprot | Total number of glycosylation sites annotated with Uniprot and detected | Total number of glycosylation sites annotated with Uniprot and not detected | Average number of glycosylation sites detected per protein | The maximum number of glycosylation sites detected per protein |
|---|---|---|---|---|---|---|---|
| *Antigen* | *2511* | *15 344* | 3,962 | 11,382 | 16,678 | 6.1 | 75 |
| *Antibody* | *136* | *142* | - | - | - | *1.0* | *3* |

**Table S2**: Detected and annotated glycosylation sites for Abs and Ags. Summary of the result section ‘Antigen Glycosylation’.

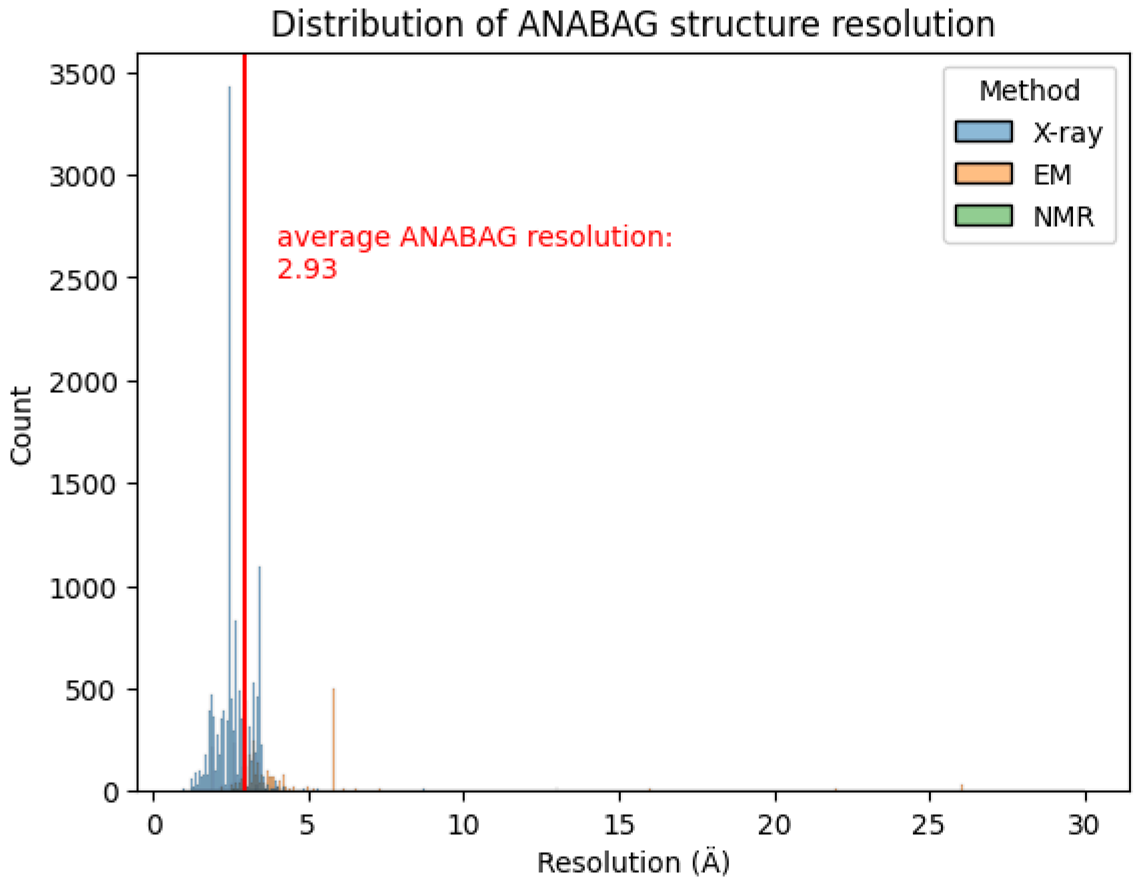


**Figure S2**: Distribution of experimental resolutions for ANABAG complexes. The average resolution of 2.93 Å is indicated in red. The different experimental methods are indicated in Blue (X-ray), Orange (Electron microscopy), and Green (Nuclear magnetic resonance)

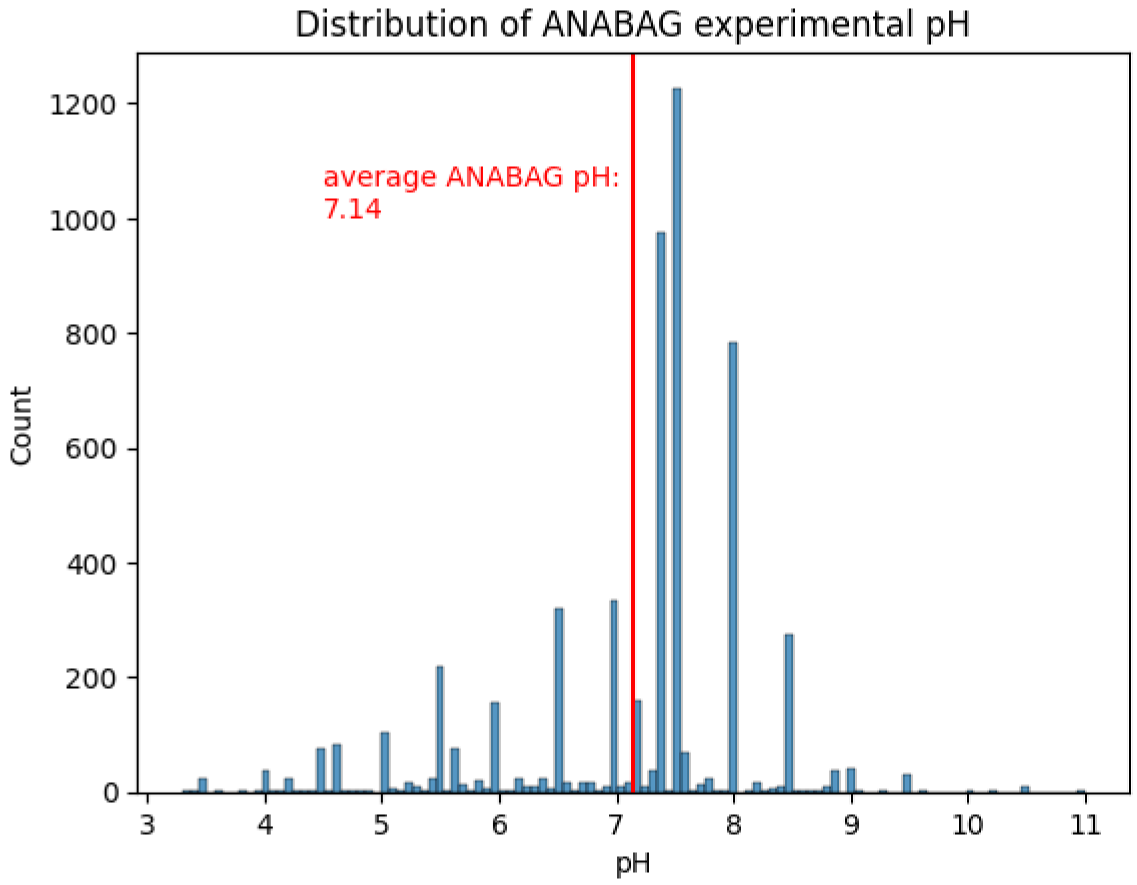


**Figure S3:** Distribution of experimental pH conditions for complexes in ANABAG. The average pH used in experiments is indicated in red (7.14), the minimum and maximum values are 3.3 and 11. The values displayed here are the last listed in the header of the PDB file.

**Additional information Figure 5**:

Alignments for paratopes of Figure 5: (epitopes are 100% identical)

*>3gbm*

TNNYSIFGSDIFSNNYYYR---

----GPFRSPIFKDDFAMGYQR

>4fqi

Identity : 26.67%

Similarity : 46.67%

**Additional information Figure 6**:

Alignments for paratopes of Figure 6: (epitopes are 100% identical)

>2vir

-------NLISNWAGNNFYDYDFY-

TTSNYGTWYSNHWAGNNYGYGVFYY

>7jti

Identity: 41.18%

Similarity : 47.06%

### Material and Methods: How to reproduce the work:

*Missing residues*:
Chains with missing regions were extracted as fasta files, and aligned with the corresponding fasta file of the PDB entry. Sequences were aligned with the Biopython pairwise aligner, and the parameters used for the alignment are the following:

```
aligner.mode = "global"
aligner.match_score = 1.0
aligner.mismatch_score = 0.0
aligner.gap_score = -0.1
aligner.target_internal_open_gap_score = -1
aligner.query_internal_open_gap_score = -1
aligner.extend_gap_score = 0
aligner.query_left_extend_gap_score = 0
aligner.target_left_extend_gap_score = 0
aligner.query_right_extend_gap_score = 0
aligner.target_right_extend_gap_score = 0
```

We used these parameters to prevent residues that were close to the gap region from being misplaced at the center of the gap, which could ultimately introduce errors in the modelling.
All missing regions were modelled in the presence of all complex chains. This prevented the modelled part from exploring conformations that would lead to structure entanglements.

*MMseq2*:
All sequences of ANABAG chains were extracted, and the Ab and Ag were separated into two Fasta file ensembles. Using these two ensembles of sequences, we used the cluster command from MMseq2 version 14-7e284, with a sensitivity of 2 and a coverage of 0. The coverage was set to 0 to prevent sequences with 100% identity from being clustered into different groups if they had different sizes. We used sequence identity thresholds of 1, 0.95, 0.8, 0.6, 0.4, 0.2 for antigens and 1, 0.95, 0.8, 0.6 for antibodies.

*Clustal-omega*:
We used Clustal-omega for building multiple sequence alignments of groups of sequences that were clustered together with MMseq. Each MMseq clustering was done using a sequence identity threshold and is referred to as a GxAG or GxAB grouping, respectively, for antigens and antibodies. The resulting groups of sequences were aligned using Clustal-omega version 1.2.4 code-name 'AndreaGiacomo' default parameters.

*Rosetta*:
Rosetta version 2021.16 release.8ee4f02 was used to score and relax the structures.
The Ag, Ab, and complex structures were minimized with the following protocol:

*relax.static.linuxgccrelease* -nstruct 1 -relax:quick. The structures used were the formatted and modeled (if there were missing regions) structures. Per residue and per structures scores were calculated with per_residue_energies.static.linuxgccrelease from the relaxed and unrelaxed structures.

*SASA*:
Residue and structure surface accessible solvent areas were calculated using freesasa version 2.0.3. Both relative accessible areas and surface accessible areas were calculated for all residues of ANABAG, and the interface as well as the protein surfaces in $A^2$ were retrieved.

*FPTS*:
Charges (in elementar units) were calculated using FPTS both in the pH condition found last in the PDB header, and at pH 7.0 with a salt concentration [NaCl] of 150mM. For each calculation, the number of equilibration steps and production steps was, respectively, 10^3 and 1.2 10^4. Charges were calculated on the separated Ag and Ab structures as well as for the whole Ab-Ag complex. The structures (formatted and modelled structures) were used as input to create the coarse-grained structures using in-house scripts. This script reads a .pdb file, extract the alpha carbon coordinates and the associated residue name. The residues are written in a .aam coarse grain file in the same order as in the PDB. At the beginning of a chain, a NTR bead is placed at the coordinates x+0.1,y+0.1,z+0.1 of the first chain residue. At the end of a chain a CTR bead is placed on the ending residue in the same way.

*Graph centrality measures, residue depth*:
Degree, betweenness centrality, eigenvector centrality, and closeness centrality for residues were calculated with the *networkx* version 2.1 package for Python. [1]
Residue depth was calculated using the biopython *residuedepth* module. The residue and Cα depth were calculated for all structure residues. The structures used for this step were the formatted and modelled structures.

*DSSP*:
Secondary structure was assigned using the *mkdssp* version 4.4.5. The entire complex was processed, and the secondary structure, φ, ψ angles were extracted from the output.

*Glycan detection*:
Glycosylated residues and glycan attached to these residues have been detected with the use of Privateer version MK V [2]. The command *privateer_exec* was used on each initial, unformatted biological unit with heteroatoms. Then the resulting files were parsed using in-house scripts to identify the glycosylations and their attached residues. The ramifications of identified glycosylations were not extracted.

[1] A. A. Hagberg, D. A. Schult, et P. J. Swart, « Exploring Network Structure, Dynamics, and Function using NetworkX », in *Proceedings of the 7th Python in Science Conference*, G. Varoquaux, T. Vaught, et J. Millman, Éd., Pasadena, CA USA, 2008, p. 11-15.
[2] J. Agirre, J. Iglesias-Fernández, C. Rovira, G. J. Davies, K. S. Wilson, et K. D. Cowtan, « Privateer: software for the conformational validation of carbohydrate structures », *Nat. Struct. Mol. Biol.*, vol. 22, nº 11, p. 833-834, nov. 2015, doi: 10.1038/nsmb.3115.
